# How much does *N*_*e*_ vary among species?

**DOI:** 10.1101/861849

**Authors:** Nicolas Galtier, Marjolaine Rousselle

**Affiliations:** ISEM, CNRS, Univ. Montpellier, IRD, EPHE, Montpellier, France; Bioinformatics Research Centre, Aarhus University, DK Aarhus, Denmark

**Keywords:** genetic drift, population size, mutation load, distribution of fitness effects, site frequency spectrum

## Abstract

Genetic drift is an important evolutionary force of strength inversely proportional to *N*_*e*_, the effective population size. The impact of drift on genome diversity and evolution is known to vary among species, but quantifying this effect is a difficult task. Here we assess the magnitude of variation in drift power among species of animals via its effect on the mutation load – which implies also inferring the distribution of fitness effects of deleterious mutations (DFE). To this aim, we analyze the non-synonymous (amino-acid changing) and synonymous (amino-acid conservative) allele frequency spectra in a large sample of metazoan species, with a focus on the primates vs. fruit flies contrast. We show that a Gamma model of the DFE is not suitable due to strong differences in estimated shape parameters among taxa, while adding a class of lethal mutations essentially solves the problem. Using the Gamma + lethal model and assuming that the mean deleterious effects of non-synonymous mutations is shared among species, we estimate that the power of drift varies by a factor of at least 500 between large-*N*_*e*_ and small-*N*_*e*_ species of animals, *i.e.*, an order of magnitude more than the among-species variation in genetic diversity. Our results are relevant to Lewontin’s paradox while further questioning the meaning of the *N*_*e*_ parameter in population genomics.

## Introduction

Genetic drift, the fluctuation of allele frequencies due to the randomness of reproduction, is one of the major evolutionary forces. Drift affects the fixation probability of selected mutations and is therefore a source of genetic load (Ohta 1972, Lynch et al. 2011). Drift impacts patterns of genome variation and can mimic or hide traces of adaptation (Jensen et al. 2005; Klopfstein et al. 2005; Peischl et al. 2018). Quantifying drift and its variation is obviously an important goal. The strength of genetic drift can be directly assessed from time series data, *i.e.*, by analysing the dynamics of allele frequency across a controlled number of generations (Jónás et al. 2016, Nené et al. 2018). This is convenient for experimentally evolving populations, but more tricky in natural conditions, where populations are less easily defined and the effects of immigration difficult to control for (Ryman et al. 2019). For these reasons, the strength of genetic drift is often approached at species level via its long-term interaction with other evolutionary forces. In a Wright-Fisher population the power of drift – i.e., the across-generation variance in allele frequency due to random sampling of organisms – is inversely proportional to the effective population size, *N*_*e*_, and issues related to the variation in drift intensity can also be phrased in terms of variation in *N*_*e*_.

The amount of neutral genetic diversity, or heterozygosity, carried by a population, π, is expected to reflect the mutation/drift balance and at equilibrium be proportional to the *N*_*e*_.μ product, where μ is the mutation rate. In principle, one could therefore assess the variation in *N*_*e*_ among species from the variation in π. Empirical evidence shows that heterozygosity is indeed correlated with abundance across species (Ellegren & Galtier 2016). The magnitude of the observed variation, however, is smaller than intuitively expected, and moderate differences in heterozygosities are sometimes reported between species that vary markedly in census population size – an observation often called Lewontin’s paradox (Lewontin 1974, Leffler et al. 2012, Romiguier et al. 2014). Three main reasons have been invoked to explain this conundrum. First, the equilibrium π is not only influenced by *N*_*e*_ but also by μ, which of course might vary between species and obscure the signal. Secondly, population size can vary in time. In this case π is expected to reflect not the contemporary *N*_*e*_, but rather the long-term *N*_*e*_, which more precisely is the time-harmonic mean of *N*_*e*_ (Wright 1938) and is strongly influenced by small values. Said differently, current genetic diversity might be largely determined by ancient bottlenecks or founder effects, irrespective of the amount of drift normally experienced by the considered population. Thirdly, selection at linked sites, either positive (Gillespie 2001) or negative (Charlesworth et al. 1995), can substantially affect π and may dominate over the effects of drift in large populations (Corbett-Detig et al. 2015, Elyashiv et al. 2016). For all these reasons, even though genomic data have confirmed the impact of drift on species genetic diversity, one cannot safely assume that π varies among species proportionally to *N*_*e*_, or in inverse proportion to the strength of drift.

Here we attempt to assess the variation in *N*_*e*_ among species by exploiting another drift-dependent variable, which is the load of segregating deleterious mutations: small-*N*_*e*_ species are expected to carry a higher load than large-*N*_*e*_ ones at selection/drift equilibrium. The segregating mutation load can conveniently be approached via the number and population frequency of non-synonymous (= amino-acid changing) variants. This implies focusing on the coding fraction of the genome, which can be seen as a limitation. On the other hand, coding sequences offer a unique opportunity to neatly control for the effect of mutation rate and demography by jointly analyzing the synonymous (= amino-acid conservative) variation, which can be assumed to be neutral. The ratio of non-synonymous to synonymous heterozygosity, π_N_/π_S_, is a typical measure of the mutation load (Chen et al. 2017). The π_N_/π_S_ ratio has a number of desirable properties. First, it is mutation rate-independent, as indicated above. Secondly, π_N_/π_S_ is expected to approach its equilibrium faster than π after a change in *N*_*e*_ (Pennings et al. 2014, Brandvain & Wright 2016, Gravel 2016). For this reason π_N_/π_S_ should be less sensitive than π to ancient bottlenecks and selective sweeps. Empirically, π_N_/π_S_ was found to be negatively correlated to population size in *Drosophila* (Jensen & Bachtrog 2011), birds (Figuet et al. 2016), animals (Romiguier et al. 2014), plants (Chen et al. 2017) and yeasts (Elyashiv et al. 2010).

So, can one quantify *N*_*e*_, or its among species variation, based on coding sequence polymorphism data? One major hurdle is that the expected amount and pattern of non-synonymous variation is determined not only by *N*_*e*_ but also by the strength of selection, or more precisely, the distribution of fitness effects (DFE) of non-synonymous mutations, which is unknown (Eyre-Walker and Keightley 2007). Drift is only expected to detectably affect the population frequency of mutations with selection coefficient, *s*, of the order of 1/*N*_*e*_ or smaller. Consider two populations of effective sizes *N*_*1*_ and *N*_*2*_. The expected difference in mutation load between the two essentially depends on the amount of deleterious mutations of effect intermediate between −1/*N*_*1*_ and −1/*N*_*2*_. If the DFE was such that most non-synonymous mutations are either much more or much less deleterious than these two values, then a small difference in load is to be expected between the two species. If however a large fraction of the non-synonymous mutations had intermediate selection coefficients, then the contrast would be higher. Welch et al. (2008) demonstrated that in a Wright-Fisher population, if the fitness effect of non-synonymous mutations follows a Gamma distribution of mean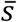and shape parameter β, then the expected π_N_/π_S_ is proportional to 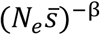. As β decreases, the distribution gets more skewed, and the expected load becomes less strongly dependent on *N*_*e*_. These considerations imply that the variation in *N*_*e*_ among species can only be assessed from the non-synonymous vs. synonymous contrast via a joint estimation of the shape of the DFE.

Eyre-Walker et al. (2006) introduced a method for estimating the DFE of non-synonymous mutations from the observed frequencies of non-synonymous and synonymous variants in a population sample – the so-called site-frequency spectra, or SFS. The idea is that slightly deleterious mutations tend to segregate at lower frequency than neutral ones, so they are expected to distort the non-synonymous SFS compared to the synonymous one. Assuming Gamma-distributed deleterious effects, expressions were derived for the expected non-synonymous and synonymous SFS at mutation/selection/drift equilibrium as a function of the population mutation rate, the shape and mean of the DFE, and nuisance parameters aimed at capturing demographic effects (Eyre-Walker et al. 2006). The method has been widely re-used since then, with modifications, mostly with the aim of estimating the adaptive amino-acid substitution rate (*e.g.* Keightley & Eyre-Walker 2007, Boyko et al. 2008, Eyre-Walker and Keightley 2009, Schneider et al. 2011, Galtier 2016, Tataru et al. 2017, Moutinho et al. 2019, Uricchio et al. 2019). The distinct versions of the method mainly differ in how they account for departures from model assumptions. These include ancient changes in *N*_*e*_ (Eyre-Walker 2002, Tataru et al. 2017, Rousselle et al. 2018a), linked selection (Messer & Petrov 2013, Uricchio et al. 2019), beneficial mutations (Galtier 2016, Tataru et al. 2017), and selfish processes such as GC-biased gene conversion, a meiotic distorter that favours G and C over A and T alleles irrespective of fitness effects. Three recent studies (Corcoran et al. 2017, Bolivar et al. 2018, Rousselle et al. 2018b) have demonstrated that GC-biased gene conversion can strongly affect inferences based on the non-synonymous vs. synonymous contrast, and must be seriously taken into account.

Using this method, one can get an estimate of the distribution of the *S*=4*N*_*e*_*s* product, and particularly its mean 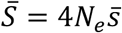. Under the assumption that distinct species share a common DFE, and therefore a common average selection coefficient 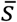 (Loewe et al. 2006), one can estimate the between-species ratio of *N*_*e*_ from the between-species ratio of 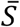. Here we analyze a recently generated population genomic data set in animals, with a focus on the primates *vs.* fruit flies comparison. We ask two questions: (i) is the DFE of non-synonymous mutations similar among species? (ii) if yes, by how much does 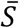, and therefore drift power, vary across species? Assuming a classical Gamma model for the DFE did not allow us to reliably assess the variation in *N*_*e*_ due to important differences in estimated shape parameter between taxa. This problem, however, was alleviated by including in the model a *N*_*e*_-independent fraction of lethal mutations. Controlling for the effect of GC-biased gene conversion and segregating beneficial mutations, we estimated that *N*_*e*_ varies by a factor of ∼500 between primates and fruit flies, *i.e*., an order of magnitude more than predicted from the variation in genetic diversity. An even wider range of variation was uncovered when we sampled more broadly across the metazoan phylogeny.

## Methods

### Data sets

We used the coding sequence polymorphism data set assembled by Rousselle et al (2019). This includes 50 species from 10 diverse taxa of animals (hereafter called “groups”), namely primates, rodents, passerines, fowls, fruit flies, butterflies, ants, mussels, earthworms and ribbon worms. Data in the former five groups (vertebrates + fruit flies) were taken from public databases. In the other five groups, which are non-model invertebrates, exon capture data were newly generated by Rousselle et al. (2019). Six to twenty individuals per species were genotyped at 531,360 to 14,112,150 coding positions from 1261 to 8604 genes (Table S1). The data set was built by selecting in each group a unique set of genes common to all species. Distinct groups, however, have distinct gene sets.

The primate and fruit fly data sets are of particularly high quality in terms of genome annotation and sample size. The two groups, furthermore, have contrasted levels of genetic diversity, with primates being among the least polymorphic, and fruit flies among the most polymorphic, taxa of animals (Leffler et al. 2012, Romiguier et al. 2014). We therefore focused on these two groups in most of the analysis. In primates, the *Papio anubis* data set had a relatively low sample size of five diploid individuals and was not analyzed here. In fruit flies, the *Drosophila sechellia* data set contained a relatively small number of SNPs and was also removed. Our main data set therefore includes five species of catarrhine primates – *Homo sapiens, Pan troglodytes, Gorilla gorilla, Pongo abelii, Macaca mulatta* – and five species of fruit flies – *D. melanogaster, D. simulans, D. teissieri, D. yakuba, D. santomea*.

In each species, the synonymous and non-synonymous SFS were generated by counting, at each biallelic position (SNPs), the number of copies of the two alleles, tri- or quadri-allelic positions being ignored. To account for missing data, a specific sample size, *n*, was chosen for each species, this number being lower than twice the number of sampled individuals. Biallelic SNPs at which a genotype had been called in less than *n*/2 individuals were discarded. When genotypes were available in exactly *n*/2 individuals, the minor allele count was simply recorded. When genotypes were available in more than *n*/2 individuals, hypergeometric projection to the {1, *n*} set was performed (Hernandez et al. 2007, Gayral et al. 2013). We used the so-called “folded” SFS in our main analysis, *i.e.*, did not rely on SNP polarization, but rather merged counts from the *k*_th_ and (*n*-*k*)_th_ categories, for every *k*. Unfolded SFS (Rousselle et al. 2019) were also used in a control analysis.

To account for the confounding effect of GC-biased gene conversion, we also generated folded GC-conservative synonymous and non-synonymous SFS by only retaining A|T and G|C SNPs, following Rousselle et al. (2018b). Estimates based on GC-conservative SFS are expected to be unaffected by any bias due to GC-biased gene conversion, but this comes at the cost of a much smaller number of SNPs. Two species, ribbonworm *Lineus longissimus* and fowl *Pavo cristatus*, had less than 100 GC-conservative SNPs and were removed from the data set.

We also re-analysed two previously-published coding sequence population genomic data sets. Chen et al. (2017) gathered SFS data from published genome-wide analyses in 34 species of animals. We focused on the 23 species in which at least five diploid individuals were sampled. These include 13 vertebrates (ten mammals, one bird, two fish), nine insects (seven *Anopheles* mosquitoes, one fruit fly, one butterfly) and one nematode. Galtier (2016) analyzed a data set of 44 species from eight distinct metazoan phyla. We selected the 28 species in which sample size was five or above, *i.e*., six species of vertebrates, six insects, five molluscs, three crustaceans, three echinoderms, two tunicates, one annelid, one cnidarian, and one nematode. All the analyzed data sets are freely available from https://zenodo.org/record/3818299#.XramS-lS88o.

### Inference methods

For each species, the population-scaled mean selection coefficient of deleterious mutations, 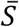, was estimated using the maximum likelihood method introduced by Eyre-Walker et al. (2006), in which a model assuming Gamma distributed deleterious effects of amino-acid changing mutations is fitted to the synonymous and non-synonymous SFS. Following up on Galtier (2016), we developed a multi-species version of this model, where the shape parameter of the Gamma distribution can be shared among species. We also implemented distinct models for the DFE, namely the Gamma+lethal, partially reflected Gamma and their combination. The Gamma+lethal model simply assumes that a fraction *p*_*lth*_ of the non-synonymous mutations are lethal, *i.e*., cannot contribute any observable polymorphism, whereas a fraction 1-*p*_*lth*_ has Gamma-distributed effects (Eyre-Walker et al. 2006, Elyashiv et al. 2010). The partially reflected Gamma model of the DFE was introduced by Piganeau and Eyre-Walker (2003) and considers the existence of back mutations from the deleterious state to the wild type. This model entails no additional parameter compared to the Gamma model, and can be easily combined with the “+ lethal” option. These methods were here newly implemented in a multi-SFS version of the grapes program (Galtier 2016, https://github.com/BioPP/grapes). The amount of neutral polymorphism of each species was assessed using the π_S_ statistics, following Romiguier et al. (2014).

### Simulations

Following Rousselle et al. (2018a), we performed simulations using SLIM V2 (Haller & Messer 2016) in order to assess how quickly π_s_ and the estimated S (Gamma model) reach their equilibrium when *N*_*e*_ varies in time. We simulated the evolution of coding sequences in a single population evolving forward in time and undergoing first an increase and then a decrease in effective population size. We considered a panmictic population of 10_4_ or 2×10_4_ diploid individuals whose genomes consisted of 1500 coding sequences, each of 999 base pairs. The mutation rate was set to 2.2×10_-7_ per base pair per generation and the recombination rate to 10_7_ per base pair per generation. The assumed distribution of the fitness effect of mutations comprised 50 % of neutral mutations and 50 % of mutations following a negative gamma distribution of mean −2.5 and shape 0.3. Each mutation that arose during a simulation was categorized as either synonymous (if the fitness effect was zero) or non-synonymous (if the fitness effect was different from zero), allowing us to compute π_n_, π_s_, and π_n_/π_s_ at any time point throughout the simulation. In order to make simulations tractable, we used small effective population sizes and high mutation rates and selection coefficients, knowing that only the products of these quantities, i.e., 4*N*_*e*_μ=0.88.10_-2_ and 4*N*_*e*_*s*=-0.025, are relevant. Simulations were replicated 50 times.

## Results

### Synonymous genetic diversity

We first analyzed synonymous and non-synonymous SFS in five species of primates and five species of fruit flies. The estimated synonymous diversity, π_S_, was much higher in fruit flies than in primates (Table 1), consistent with the literature (Leffler et al. 2012). The estimated π_S_ varied from 0.062% in *H. sapiens* to 3.2% in *D. yakuba, i.e.*, a factor of 51 (figure 1, X-axis). The per group median π_S_ was 0.11% in primates and 1.5% in fruit flies, *i.e.*, varied by a factor of 13.

**Table 1.** Maximum likelihood estimates of 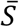 in primates and fruit flies under the Gamma+lethal DFE model.

| species | $\pi_s(\%)$ | $\beta$ shared across species | | one $\beta$ per group | |
| --- | --- | --- | --- | --- | --- |
| | | $p_{lth}=0.6$ | $p_{lth}=0.65$ | $p_{lth}=0.6$ | $p_{lth}=0.65$ |
| *H. sapiens* | 0.06 | -3.36 | -1.39 | -2.75 | -1.53 |
| *G. gorilla* | 0.11 | -4.38 | -2.08 | -4.18 | -2.22 |
| *P. troglodytes* | 0.11 | -7.56 | -3.30 | -7.03 | -3.45 |
| *P. abelii* | 0.16 | -9.16 | -4.64 | -9.64 | -4.59 |
| *M. mulatta* | 0.18 | -20.0 | -10.4 | -24.5 | -9.64 |
| *D. santomea* | 0.11 | -401 | -232 | -395 | -234 |
| *D. melanogaster* | 1.07 | -502 | -295 | -493 | -297 |
| *D. teissieri* | 1.52 | -1860 | -1080 | -1820 | -1090 |
| *D. yakuba* | 1.57 | -946 | -553 | -928 | -558 |
| *D. simulans* | 3.18 | -1660 | -970 | -1630 | -979 |
| **max/min $N_e$** | **53.0** | **555**<br>[324-1050] | **779**<br>[438-1660] | 662<br>[353-1430] | 714<br>[422-1350] |
| **group-median $N_e$ ratio** | **13.8** | **125**<br>[84-203] | **168**<br>[102-296] | 132<br>[87-237] | 162<br>[103-281] |

**Figure 1:**
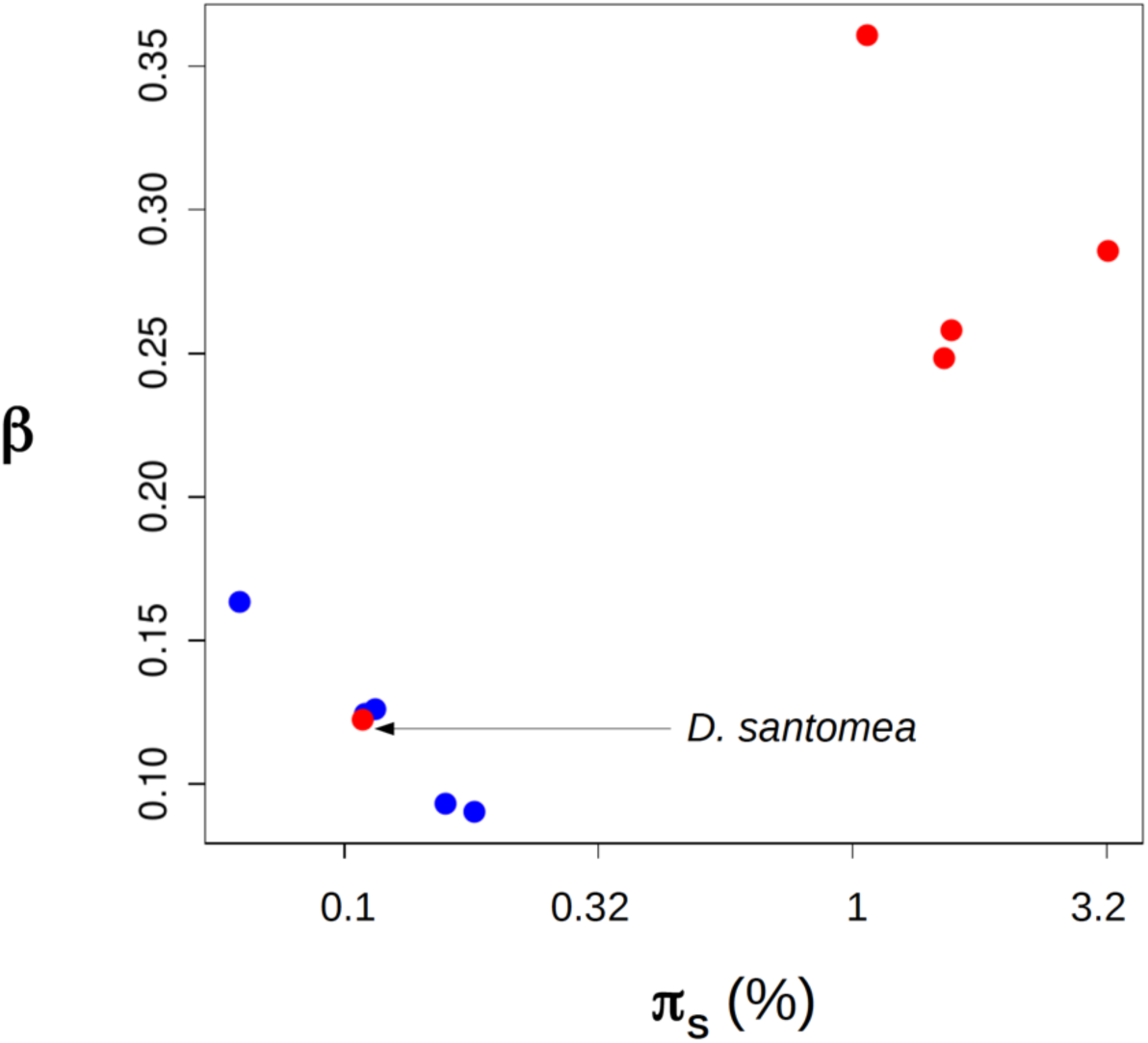
The estimated Gamma shape differs between primates and fruit flies. Blue dots: primates. Red dots: fruit flies. X-axis: synonymous genetic diversity (log_10_ scale). Y-axis: estimated shape parameter assuming a Gamma DFE of non-synonymous mutations.

### Gamma DFE

A model assuming Gamma-distributed population scaled selection coefficient was fitted to SFS data separately in the 10 species. We uncovered substantial variation in β, the shape parameter of the Gamma DFE: the maximum likelihood estimate of β varied from 0.09 to 0.36 among species (figure 1, Y-axis). When we instead fitted a model assuming a common β across species, the likelihood dropped severely (one β per species, log-likelihood=-1100.8; shared β, log-likelihood=-1382.7), and the hypothesis of a common DFE shape across the 10 species was strongly rejected by a likelihood ratio test (LRT; *p*-val<10_-20_; nine degrees of freedom). There was a trend for species from the same group to provide similar estimates of β (figure 1). In primates, the range of estimated β was narrow ([0.09;0.16]) and the hypothesis of a common DFE shape was not rejected by LRT (optimal β for primates: 0.11; *p*-val=0.45; four degrees of freedom). In fruit flies, the optimal β was close to 0.3 in four species, but much lower in *D. santomea* (figure 1). The optimal fruit fly β was firmly rejected by primates as a group (LRT; *p*-val<10_-20_; one degree of freedom), and reciprocally (LRT; *p*-val<10_-20_; one degree of freedom; the test includes *D. santomea*).

Estimates of 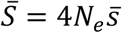, the population scaled mean selection coefficient, varied greatly among species in this analysis but this was largely explained by variations in β. Gamma shape and mean are known to be correlated parameters, and this was verified here: the across species correlation coefficient between log-transformed 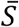 and log-transformed β was −0.94 in primates and −0.99 in fruit flies (figure S1). The among-species variation in estimated 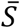 can therefore not be taken as a reliable indicator of the variation in *N*_*e*_ when β differs much among species.

The observed variation in estimated DFE shape might in principle reflect biological differences among species and groups, *e.g.*, differences in composition/structure of the proteome/interactome, maybe strengthened by our gene sampling strategy – here, species from the same group share the same genes, whereas distinct groups have distinct gene sets. The *D. santomea* behavior, however, appears difficult to reconcile with this hypothesis. *D. santomea* is closely related to *D. yakuba* and *D. teissieri* (Turissini and Matute 2017), and shares the same gene set as other species of fruit flies. We see no obvious reason why the DFE of deleterious non-synonymous mutations in *D. santomea* would differ strongly from other fruit flies, and resemble the primate DFE. Interestingly, *D. santomea* shares with primates a relatively low genetic diversity (figure 1), perhaps as a consequence of its restricted geographic distribution (Bachtrog et al. 2006). For this reason, we hypothesized that the among-species variation in estimated β we report could reflect a failure of the Gamma model to capture the details of the DFE at all values of *N*_*e*_, rather than genuine differences in DFE among species.

### Gamma + lethal DFE

It should be recalled that very highly deleterious alleles have essentially zero probability of being observed at polymorphic stage with the sample size we used here. We reasoned that, if the DFE included a proportion of mutations of effects essentially independent of *N*_*e*_, this could lead to undesired effects when fitting a Gamma distribution of *S*, a variable proportional to *N*_*e*_.

To investigate this, we fitted to SFS data a model assuming a proportion *p*_*lth*_ of lethal mutations, and a proportion (1-*p*_*lth*_) of mutations of Gamma distributed effects. Lethal mutations are here defined as mutations having a probability of being observed at polymorphic stage equal to zero. Figure 2 shows how the likelihood responds to *p*_*lth*_ and β in primates and fruit flies. In fruit flies, the optimal β was close to 0.3 irrespective of *p*_*lth*_, and the likelihood was maximal when *p*_*lth*_ was close to 0.75. In primates, the optimal β varied more visibly with *p*_*lth*_. It was close to 0.1 when *p*_*lth*_ was low, as indicated above, but increased towards higher values when *p*_*lth*_ increased. The maximal likelihood in primates was still obtained when *p*_*lth*_ was close to zero and β close to 0.1, but importantly, areas of the parameter space close to the fruit fly optimum (*e.g., p_lth_*∼0.65 and β∼0.3) provided a reasonably good fit to the data (figure 2). This suggests that the DFE perhaps does not differ so dramatically between primates and fruit flies, offering the opportunity to compare estimates of 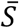 across species under the Gamma+lethal model. The difference in log-likelihood between the one-β-per-species and shared-β models was substantially decreased under the Gamma+lethal model (191.0) compared to the Gamma model (281.9). Still, the one-β-per-species model significantly rejected the shared-β model in both cases, indicating that the inclusion of the *p*_*lth*_ parameter was not sufficient to erase every perceptible difference in DFE among species.

**Figure 2:**
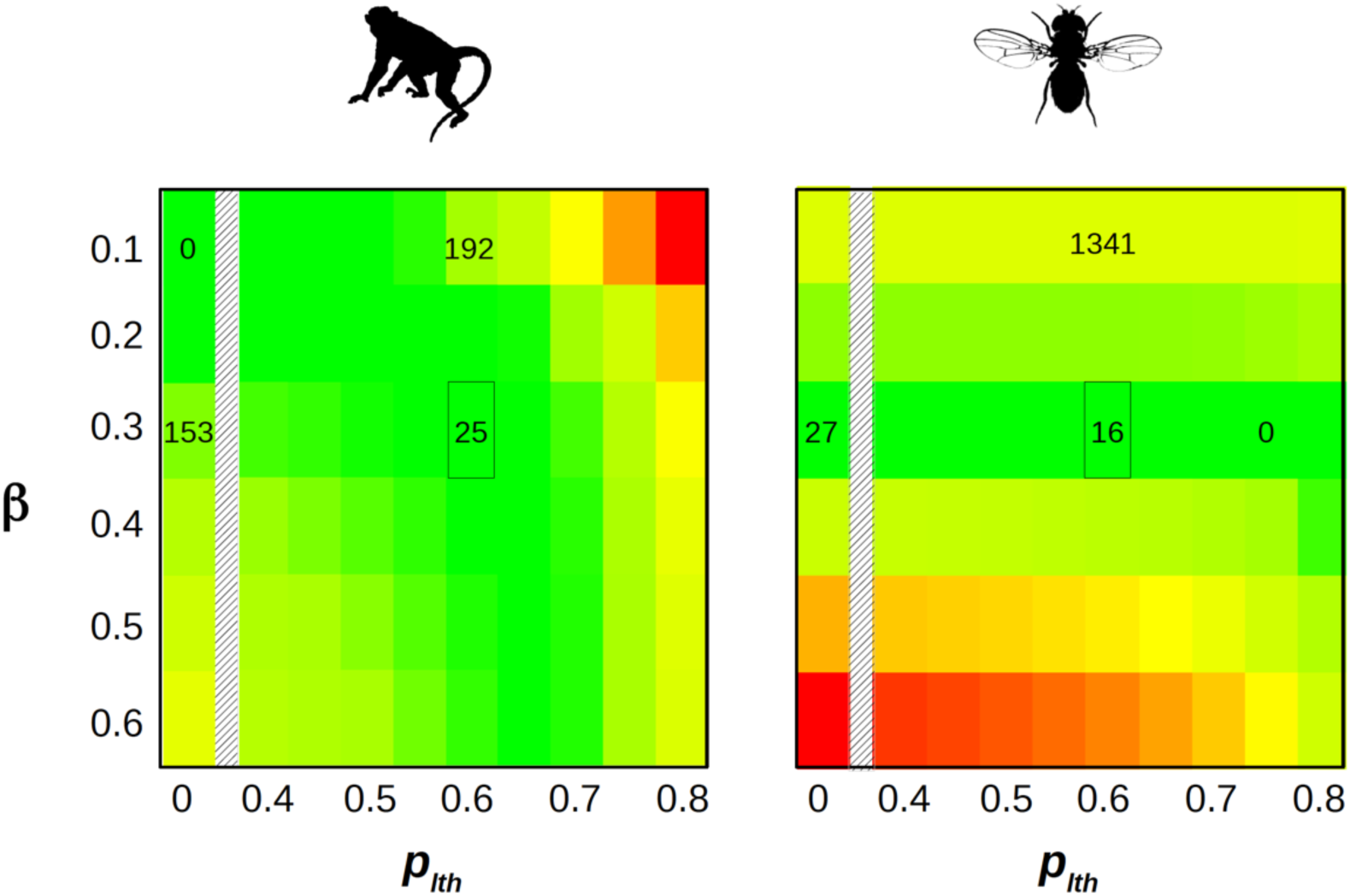
Gamma+lethal model, likelihood surface. X-axis: proportion of lethal non-synonymous mutations. Y-axis: Gamma shape parameter for non-lethal, non-synonymous mutations. The color scale is indicative of the log-likelihood: green=high, yellow=intermediate, red=low. The difference between local and maximal log-likelihood is indicated by numbers within a couple of relevant cells – 0 means maximal. The marked cell corresponds to the maximum of the likelihood when both data sets are jointly analyzed (see Figure S2). X-axis continuously covers the 0.4-0.8 range, and also shows the *p*_*lth*_=0 case (Gamma model).

We jointly analyzed the ten species from the two groups under the Gamma+lethal model assuming a shared shape parameter among species, and found that the likelihood was maximal when *p*_*lth*_ was in the range 0.6 – 0.65 (figure S2). We considered these two values as plausible estimates of *p*_*lth*_. When *p*_*lth*_ was set to 0.6, the maximum likelihood estimate of β was 0.278. In this analysis the maximum likelihood estimate of 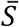 varied by a factor of 550 among species, and the median fruit fly 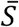 was 125 times as large as the median primate 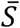 (Table 1). Of note, the estimated 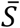 in *D. santomea* was similar to that of other species of fruit flies, and much larger than in primates. Setting *p*_*lth*_ to 0.65 instead of 0.6 yielded similar results (Table 1). Confidence intervals were obtained by bootstrapping SNPs (100 replicates). The Gamma+lethal model outperformed the Gamma model in this analysis: letting *p*_*lth*_ be different from zero increased the log-likelihood by 85 units, which was highly significant (*p*-val<10_-20_, one degree of freedom). We also estimated 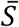 under the Gamma + lethal model assuming that the β parameter was shared by species within a group, but could vary between the two groups. Estimates of 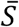 and *N*_*e*_ ratios were close to the ones obtained assuming shared β across all species (Table 1, right-most two columns), confirming that the inferred DFEs of the two groups tend to converge when a fraction of lethal mutations are included. We checked that our estimate of 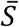 and β were such that the mean π_n_/π_s_ was linearly related to 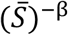 across species, as expected at equilibrium with Gamma distributed fitness effects (Welch et al. 2008). This was the case under both the Gamma and Gamma+lethal models (figure S3).

### Robustness of the estimates

We performed a number of additional analyses in order to investigate the robustness of the above results to various methodological settings. The results are summarized in Table 2. In each of the four control analyses – “GC-conservative”, “reflected Gamma”, “unfolded”, “subsampled fruit flies” – we tried eight values of *p*_*lth*_, from 0.4 to 0.75. We optimized β and the other parameters conditional on *p*_*lth*_ and recorded the likelihood. Table 2 shows the results for values of *p*_*lth*_ within 2 log-likelihood units of its maximum.

**Table 2.**
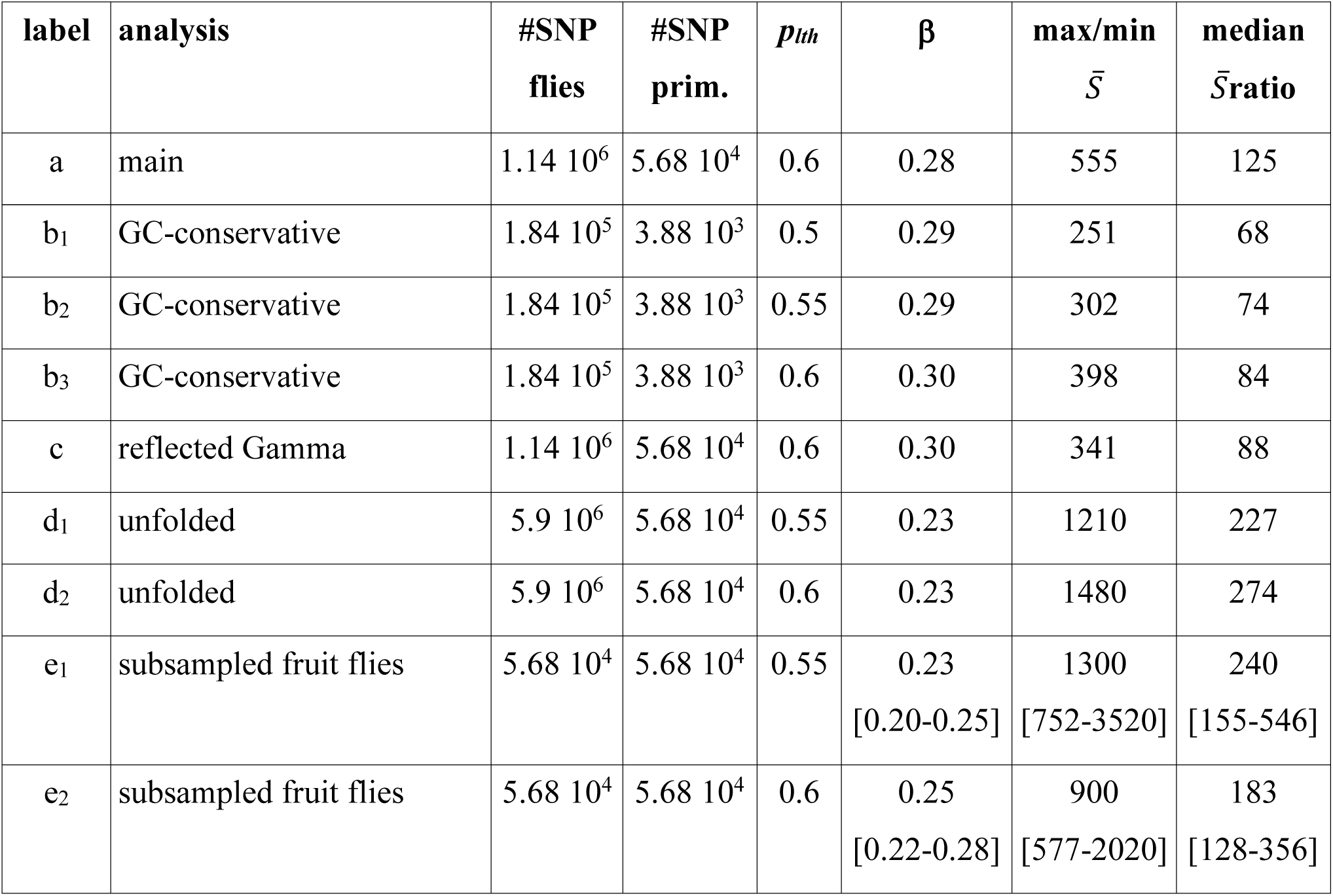
Robustness analysis, primates vs. fruit flies.

| label | analysis | #SNP<br>flies | #SNP<br>prim. | $p_{lth}$ | $\beta$ | max/min<br>$\bar{S}$ | median<br>$\bar{S}_{ratio}$ |
| --- | --- | --- | --- | --- | --- | --- | --- |
| a | main | $1.14 \cdot 10^6$ | $5.68 \cdot 10^4$ | 0.6 | 0.28 | 555 | 125 |
| b<sub>1</sub> | GC-conservative | $1.84 \cdot 10^5$ | $3.88 \cdot 10^3$ | 0.5 | 0.29 | 251 | 68 |
| b<sub>2</sub> | GC-conservative | $1.84 \cdot 10^5$ | $3.88 \cdot 10^3$ | 0.55 | 0.29 | 302 | 74 |
| b<sub>3</sub> | GC-conservative | $1.84 \cdot 10^5$ | $3.88 \cdot 10^3$ | 0.6 | 0.30 | 398 | 84 |
| c | reflected Gamma | $1.14 \cdot 10^6$ | $5.68 \cdot 10^4$ | 0.6 | 0.30 | 341 | 88 |
| d<sub>1</sub> | unfolded | $5.9 \cdot 10^6$ | $5.68 \cdot 10^4$ | 0.55 | 0.23 | 1210 | 227 |
| d<sub>2</sub> | unfolded | $5.9 \cdot 10^6$ | $5.68 \cdot 10^4$ | 0.6 | 0.23 | 1480 | 274 |
| e<sub>1</sub> | subsampled fruit flies | $5.68 \cdot 10^4$ | $5.68 \cdot 10^4$ | 0.55 | 0.23<br>[0.20-0.25] | 1300<br>[752-3520] | 240<br>[155-546] |
| e<sub>2</sub> | subsampled fruit flies | $5.68 \cdot 10^4$ | $5.68 \cdot 10^4$ | 0.6 | 0.25<br>[0.22-0.28] | 900<br>[577-2020] | 183<br>[128-356] |

The first line of Table 2 recalls some of the results presented in Table 1. In the “GC-conservative” control, we applied the same method as in the main analysis but only including C/G and A/T SNPs, which are supposedly immune from GC-biased gene conversion. The “reflected Gamma” analysis used the same data set as the main one, but assumed a reflected Gamma instead of a Gamma DFE, thus accounting for the presence of slightly beneficial mutations – in addition to a proportion of lethal mutations. The “unfolded” control used unfolded instead of folded SFS. Finally, in the “subsampled fruit flies” control, we reduced the size of the fruit fly data set to the size of the primate one, such that the two groups have equal weights in the analysis. A hundred data sets were generated by randomly subsampling in *D. melanogaster* the number of SNPs available in *G. gorilla*, and similarly for *D. santomea*/*H. sapiens, D. simulans*/*M. mulatta, D. teissieri*/*P. troglodytes*, and *D. yakuba*/*P. abelii*. The most likely value for *p*_*lth*_ was 0.6 in 77 subsampled data sets and 0.55 in the other 23. Table 2 reports the median estimates and 95% confidence intervals across the 100 subsampled data sets for these two values of *p*_*lth*_.

The GC-conservative data set included roughly ten times less SNPs than the main one. The likelihood surface was flatter and close to its maximum at three values of *p*_*lth*_ – 0.5, 0.55, 0.6 – while the estimated β was always close to 0.3. The ratio of 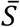 between large-*N*_*e*_ fruit flies and small-*N*_*e*_ primates was roughly twice as low as in the main analysis, either using extreme estimates or within-group medians. The reflected Gamma analysis also yielded ratios of 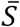 a bit lower than the main analysis – but still substantially higher than ratios of π_S_, see Table 1. Of note, the maximum log-likelihood under the reflected Gamma + lethal model (−1301.6) was slightly decreased compared to Gamma + lethal (−1297.7). When unfolded SFS were analyzed, the estimated β was lower than in the main analysis, and the ratios of 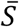 was twice as large. A similar pattern was obtained when we subsampled the fruit fly data set so as to match the primate sample size – a slightly lower estimate of β, and a max/min *N*_*e*_ ratio of the order of 10_3_. In all these control analyses the Gamma+lethal model fitted the data significantly better than the Gamma model.

### Additional metazoan taxa

We similarly analyzed an additional 37 species from eight diverse groups of animals (Rousselle et al. 2019). In this 47-species data set, the estimated synonymous diversity π_S_ varied from 0.062% in *H. sapiens* to 4.4% in mussel *Mytilus trossulus*. The within-group median π_S_ varied from 0.012 in primates to 3.3% in mussels, *i.e.*, by a factor of 27.

The results were largely consistent with the primates vs. fruit flies comparison. First applying the Gamma model, we detected a strong group effect on the estimated β (figure S4). Butterflies and ants, for instance, tended to yield relatively high estimates of β, whereas the best fit in primates and birds was reached at relatively low β. This was mitigated by moving to the Gamma+lethal model, even though the specificity of particular groups was still apparent (figure S5). When we jointly analyzed all species assuming a Gamma+lethal DFE using folded SFS and all mutations, the maximal likelihood was reached when *p*_*lth*_ was 0.5 (with *p*_*lth*_=0.55 being close) and the estimated β was 0.26. We calculated the within-group median 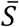, which varied by a factor of 110 among groups (Table 3, first colon). We performed the same GC-conservative, reflected Gamma and unfolded analyses as described above (Table 3). The estimated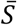in the reflected Gamma, unfolded and main analyses were strongly correlated (r_2_>0.97 in all three pairwise comparisons), while the ratio of max/min median 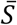 varied a bit: it was 73 in the Gamma reflected analysis, 200 in the unfolded analysis. In all three analyses, fruit flies were the group with the highest median 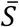, and primates the lowest.

**Table 3.** Analysis of 47 species in 10 groups of animals.

|  | main | GC-cons. | reflected | unfolded |  |
| --- | --- | --- | --- | --- | --- |
| $p_{lth}$ | 0.5 | 0.55 | 0.5 | 0.45 | |
| $\beta$ | 0.26 | 0.28 | 0.29 | 0.22 | |
| **relative median $\bar{S}$:** |  |  |  |  | **color :** |
| *primates* | 1 | 1.6 | 1 | 1 | blue |
| *ants* | 1.6 | 1 | 1.6 | 1.6 | yellow |
| *butterflies* | 1.8 | 6.1 | 1.7 | 1.8 | orange |
| *passerines* | 3.1 | 4 | 2.9 | 3.1 | magenta |
| *earthworms* | 4.8 | 6.4 | 4.3 | 5.1 | cyan |
| *rodents* | 5.2 | 7.6 | 4.6 | 6.5 | grey |
| *fowls* | 21 | 7.8 | 17 | 25 | green |
| *ribbonworms* | 38 | 120 | 29 | 55 | black |
| *mussels* | 40 | **180** | 30 | 60 | brown |
| *fruit flies* | **110** | 130 | **75** | **200** | red |

The GC-conservative analysis differed a bit and did not rank the ten groups in the same order (Table 3). In this analysis, the lowest median 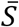 was found in ants and the highest in mussels, the ratio between these two numbers being 180. The GC-conservative estimates of 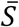 were more strongly correlated with species propagule size, a variable previously identified as a proxy for *N*_*e*_ in animals (Romiguier et al. 2014), than were the other three estimates (GC-conservative: r_2_=0.27, *p*-val=3.10_-4_; main analysis: r_2_=0.11, *p*-val=3.10_-2_; log-transformed variables). Figure 3 shows the relationship between π_S_ and the relative 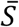 as estimated from the GC-conservative data set. Note that the Y-axis encompasses three orders of magnitude, vs. two on the X-axis. As mentioned above, *D. santomea* was an outlier: π_S_ in this species was low, but the estimated 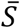 was typical of fruit flies and other large-*N*_*e*_ species. The ratio of max/min estimated 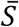 across the 47 species was 491, 1100, 301 and 1076, respectively, in the main, GC-conservative, reflected Gamma and unfolded analyses.

**Figure 3:**
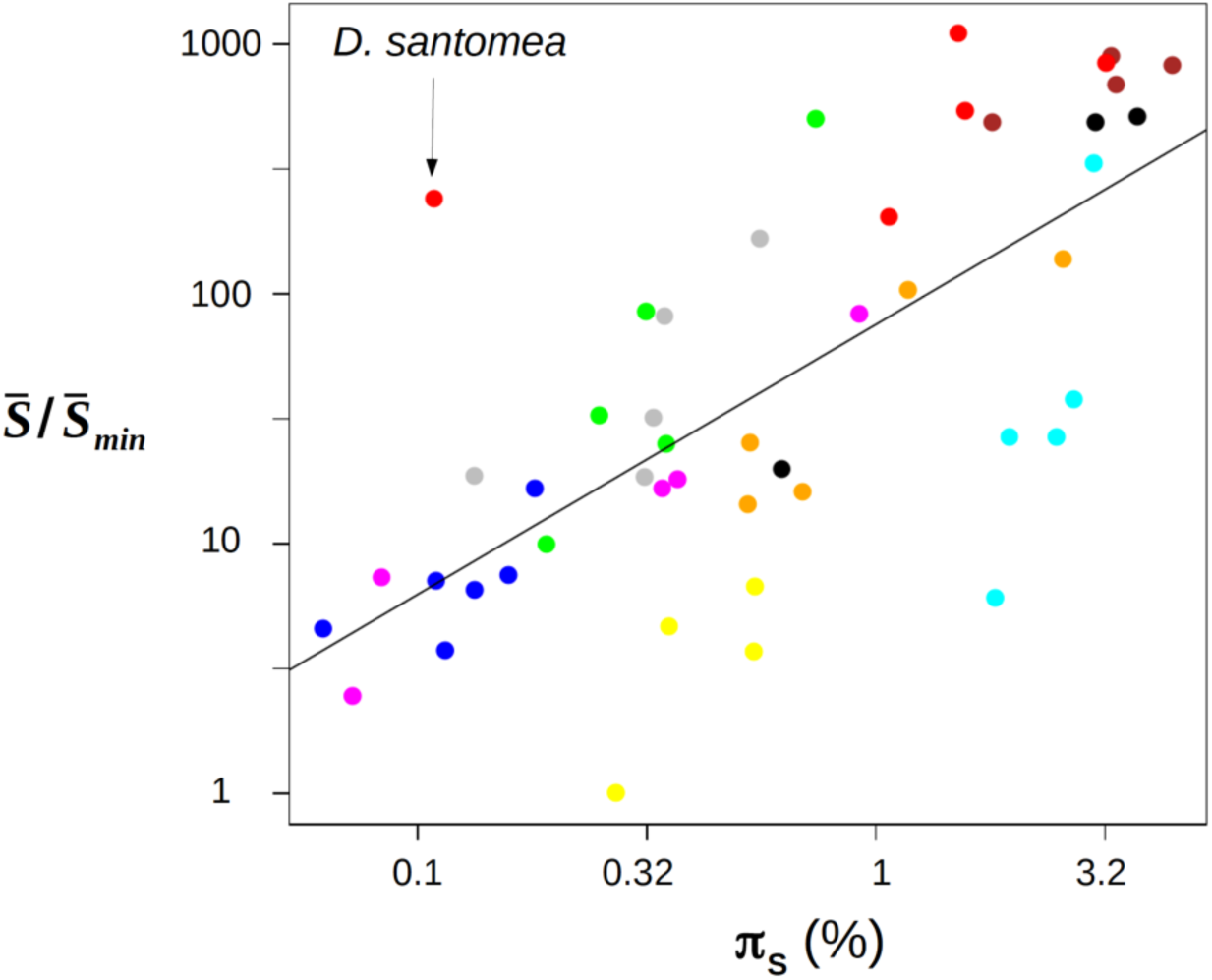
Magnitude of variation in population scaled mean selection coefficient vs. neutral genetic diversity across 47 species of animals. Each dot is for a species of animals. X-axis: relative synonymous heterozygosity. Y-axis: relative estimated 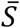. Colors indicate groups, see Table 3. Estimates of 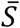 were divided by its minimal value, which was obtained in *Formica pratensis.*

We similarly analyzed two additional, recently published data sets. The Chen et al. (2017) data set (23 species) yielded results similar to the Rousselle data set: the optimal *p*_*lth*_ for this data set was 0.5, the estimated β was 0.29, and the ratio of maximal to minimal 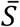 was 581 – to be compared to a max/min ratio of π_S_ of 54. Of note, the estimated ratio of 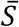 between *D. melanogaster* and *H. sapiens* was 139 in the Chen et al. (2017) data set, which is close to the 119 obtained in our main analysis (see Table 1). The Galtier (2016) data set (28 species) differed from the other two in that the likelihood was largely insensitive to *p*_*lth*_: it varied by less than 2 log units over the 0-0.6 range for *p*_*lth*_. The estimated β was close to 0.275 irrespective of *p*_*lth*_ (in the 0-0.6 range), and the ratio of maximal to minimal estimated 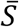 was 7300-7900, *i.e*., an additional order of magnitude compared to Rousselle’s and Chen’s data sets. This result was mainly explained by a very high estimated 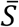 in the mosquito *Culex pipiens* and, particularly, the nematode *Caenorhabditis brenneri* (figure S6).

## Discussion

Here we introduced a novel approach to compare the intensity of genetic drift among species based on coding sequence SFS data. Below we discuss the assumptions, merits and limitations of this approach (subsection 1 to 3), before moving to the interpretation and implications of our results (subsection 4 and 5).

### 1. Estimating N_e_-related parameters from SFS data

Here we used the approach introduced by Eyre-Walker et al. (2006) for fitting a population genetic model to a synonymous and a non-synonymous SFS. This model includes three parameters of interest: population mutation rate θ, average deleterious effect 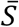 and DFE shape β. In addition, the model has *n*/2 nuisance parameters, where *n* is the sample size – the so-called *r*_*i*_’s (see equations 1 and 2 in Eyre-Walker et al. 2006, here applied to folded SFS). Parameter *r*_*i*_ multiplies the *i*^th^ entry of the expected synonymous and non-synonymous SFS. The *r*_*i*_’s are intended to capture any effect that similarly affects the fate of synonymous and non-synonymous mutations – such as linked selection, population substructure and departure from demographic equilibrium. In practice *r*_*1*_ is set to one, and *r*_*i*_, *i*>1, can be interpreted as the relative effective mutation rate of the *i*^th^ frequency category, compared to the first category.

Including the *r*_*i*_’s in the model is often necessary in terms of goodness of fit. This was the case here: when we set all the *r*_*i*_’s to 1, *i.e.*, assumed panmixy, no linked selection and demographic equilibrium, the likelihood dropped dramatically, from −1297.7 to −71,404.7 (primates + fruit flies dataset, Gamma+lethal, *p*_*lth*_=0.6). Such a simplistic model is strongly rejected by the data and can hardly be used for inference purposes. Alternatively, one could try to explicitly model population substructure, linked selection (Good et al. 2014) and/or departure from demographic equilibrium (Evans et al. 2007, Keightley & Eyre-Walker 2007). In practice, however, these effects are very difficult to disentangle. Messer and Petrov (2013), for instance, demonstrated that linked selection severely confounds inferences on the variation of *N*_*e*_ in a two-epoch model. This is why the flexible *r*_*i*_-based parametrization has been used in numerous recent applications of the extended McDonald-Kreitman method (e.g. Galtier 2016, Tataru et al. 2017, Rousselle et al. 2018, Moutinho et al. 2019).

Introducing the *r*_*i*_’s, however, has one drawback, which is that this tends to blur the interpretation of the estimates of parameters of interest, particularly θ and 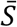. Consider for instance a data set in which the estimated *r*_*i*_’s (*i*>1) are all well below one, meaning that singletons are in excess compared to the standard coalescent expectation. Such a pattern is expected for a population having experienced a recent, strong bottleneck. What exactly θ measures in this case is unclear, in the absence of a formal model. Now consider another data set yielding the same estimate of θ, but with estimated *r*_*i*_’s (*i*>1) all well above one, as expected under gradual population decline. To conclude that the two considered populations have the same population mutation rate would appear somehow meaningless. A similar problem possibly applies to the interpretation and among-data sets comparisons of the estimated 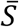. In this study we did not estimate θ via the maximum likelihood method, but rather used π_S_ as our estimate of θ. We did, however, use the maximum likelihood of 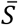. One should keep in mind that the meaning of these estimates are somehow dependent on the estimated *r*_*i*_’s, to an extent currently difficult to quantify in the absence of a formal investigation of this issue.

To investigate this issue a bit deeper, we plotted the estimated *r*_*i*_’s for all SNP categories *i* in primates and fruit flies (figure S7). In this analysis, the Gamma+lethal model was used, species were analyzed separately (no shared parameter), and *p*_*lth*_ was set to 0.6. Figure S7 shows that the vast majority of the *r*_*i*_’s belong to the [0.5, 1.5] interval, *i.e.*, do not dramatically differ from 1, both in primates and fruit flies. The figure also shows that, although the median differs a bit between primates and fruit flies for SNP category 3 to 8, the distributions are largely overlapping among the two groups. Although a more detailed analysis would be worthwhile, figure S7 does not suggest to us that the *r*_*i*_’s pose a major problem of comparability in this analysis.

### 2. Strong variation in Gamma-DFE shape across species

When a Gamma DFE was assumed, our analysis revealed significant among-taxa variation in the estimated shape parameter. In primates, the best fit was achieved when the shape parameter was of the order of 0.1 – 0.15, and consistently so in five different species. These values are close to those obtained by Castellano et al. (2019), who similarly analyzed coding sequence SFS data in nine populations of great apes. Assuming a Gamma DFE, these authors found that the best model was one in which the shape parameter was shared among species and equal to 0.16. This estimate was only slightly increased when a fraction of beneficial mutations was modeled (Castellano et al. 2019). In *Drosophila*, in contrast, we found that the value of β that best fitted the data was close to 0.3 – with the exception of *D. santomea*. This also is consistent with previous reports. Keightley et al. (2016), for instance, obtained a point estimate of 0.35 for β in *D. melanogaster*, taking special care of the problem of SNP mis-orientation. In butterflies, our median estimated β, 0.37 (min=0.29, max=0.52), is close to the 0.44 reported by Mackintosh et al. (2019) based on a different approach. Our results corroborate those of Chen et al (2017), who provided a detailed analysis of the variation in estimated β across various taxa of animals and plants, and like us reported a tendency for species with a low genetic diversity to exhibit low values of β.

The variation in estimated β we and Chen et al (2017) report is substantial. It should be recalled that the shape parameter has a strong effect on the Gamma distribution, as it is inversely proportional to its variance. Consider for example Gamma-distributed DFEs sharing the same mean *s* of, say, 0.1, but with distinct shape parameters. If β is set to 0.1, as estimated in primates, then 53% of mutations are associated with a selection coefficient smaller than 0.001. If β was rather equal to 0.3, as in fruit flies, then this percentage would be 19%, and down to 12% if β=0.4, as in butterflies. Given the scarcity of experimental data on the non-synonymous DFE in animals, one cannot firmly argue that such differences are implausible. We note, however, that if the estimates of β we obtained reflected a biological reality, this would entail considerable variation in the prevalence of small effect mutations across animal proteomes, for which an explanation would be needed.

Here we rather hypothesized that the among-taxa variation in estimated β is, for its largest part, due to model mis-specification, that is, we suggest that the true DFE might not be Gamma-distributed. Besides the above intuitive argument, this hypothesis is based on two observations. The first one is the behavior of the *D. santomea* data set, which carries much less diversity than other species of fruit flies, and yielded a distinctively lower estimate of β, suggesting that the Gamma model fails to correctly capture the shape of the DFE at all values of *N*_*e*_. The alternative explanation, *i.e.*, that the DFE in *D. santomea* truly differs from that of other fruit flies, appears dubious in this case given the low between-species divergence. Secondly, we found that the Gamma + lethal model provided a significantly better fit to the data than the Gamma model, while predicting DFE shapes that were more similar across taxa.

In this study, therefore, we implicitly attributed the observed variation in estimated Gamma shape to a methodological artifact. This is obviously questionable. It could be that the among-taxa variation in estimated β reported here and in Chen et al. (2017) has some biological relevance. This would deserve to be investigated experimentally, *e.g.* following the approach of Böndel et al. (2019), who obtained an estimate of β=0.3 in green algae *Chlamydomonas reinhardtii* based on crosses and fitness measurements in mutation accumulation lines. It should be kept in mind that our discussion of the variation in *N*_*e*_ is conditional on the assumption of a common DFE among species.

### 3. Modeling lethal mutations

The Gamma + lethal model, which considers an *N*_*e*_-independent fraction of large-effect deleterious mutations, provided a significantly better fit to the data than the Gamma model. Here is a possible interpretation of this result. Fitting a DFE model to a non-synonymous and a synonymous SFS implies accommodating both the difference in shape (the relative frequencies of singletons, doubletons, etc…) and in size (the total number of SNPs) between the two spectra. The former is determined by the balance between neutral and slightly deleterious mutations, whereas the latter mainly reflects the proportion of strongly deleterious mutations. We suggest that the Gamma distribution struggles to accommodate these two aspects at the same time. When the π_N_/π_S_ ratio is relatively high, as in primates, there is a tendency for the Gamma model to converge towards low values of β, thus ensuring a large proportion of small effect non-synonymous mutations, while a higher β could fit the difference in shape between the two spectra equally well, or maybe better. The additional *p*_*lth*_ parameter of the Gamma + lethal model, we suspect, somehow releases this constraint by controlling to a large extent the predicted π_N_/π_S_ ratio.

This interpretation seems to fit reasonably well our analysis of the primates + fruit flies dataset, and the report by Chen et al (2017) of a lower estimate of β in small-*N*_*e*_ species. This rule, however, does not always apply. Our estimate of β in *Formica* ants, for instance, was close to 0.4 under the Gamma model, despite a low genetic diversity in this taxon. More work would appear needed to determine whether the relatively high estimate of β we obtained in ants reveals a real peculiarity of this group, or is due to the specific gene set we analyzed here, or can be explained by the relatively small SNP sample size of our ant data set, compared to primates and fruit flies (Table S1). To our knowledge, the Gamma + lethal model has been tried in two studies before this one. Eyre-Walker et al. (2006) analyzed a data set of 320 genes in 90 human individuals, and found that adding the *p*_*lth*_ parameters did not change the picture much, compared to the Gamma model. This contrasts with our analysis and highlights the sensitiveness of this kind of analyses to the specificities of the data – number of genes, of individuals, SNP calling procedure. Elyashiv et al. (2010) applied the Gamma + lethal model to yeast data and found that the shape parameters converged towards 0.35 under this model, which is very close to our joint estimate, while the Gamma model supported a higher value for β.

The Gamma + lethal model is a simple modification of the Gamma model, which was sufficient to significantly improve the fit in this analysis. The model, however, is not entirely satisfactory. In particular, our analysis makes the assumption of a common proportion of very strongly deleterious mutations among species with different *N*_*e*_, which appears awkward knowing that the probability for a mutation to segregate at observable frequency is determined by the *N*_*e*_.*s* product. There might be more efficient, continuous ways to model the DFE and solve the problem posed by the Gamma distribution with this data set. One intrinsic difficulty with SFS model fitting is that deleterious mutations of sufficiently large effects will be equally unobservable irrespective of their precise selection coefficient – for instance, mutations of selection coefficient *s*=-100/*N*_*e*_ or *s*=-1000/*N*_*e*_ just have essentially zero probability to be observed in a population sample. This means that we lack information on the tail of the distribution we are trying to model.

### 4. Estimating N_e_: mutation load vs. diversity

Our analysis of the load of deleterious mutations uncovered three (based on the Rousselle et al. 2019 and Chen et al. 2017 data sets) or four (based on the Galtier 2016 data set) orders of magnitude of variation in drift power among species of animals, when the neutral genetic diversity of the very same species varied by a factor of 100 or less. Below we discuss potential reasons for this discrepancy.

First, it should be recalled that, unlike the non-synonymous to synonymous contrast, the genetic diversity of a species is influenced by the mutation rate. If the mutation rate was negatively correlated with effective population size, and differed by one or two orders of magnitude between species of animals, then our results could be explained very simply. Empirical estimates in humans (Kong et al. 2012) and *Drosophila* (Keightley et al. 2009) indeed seem to point to an order of magnitude of difference in per base, per generation mutation rate between these two taxa. We lack, however, a reliable estimate of the mutation rate in the vast majority of the species of our data set. The existence of a negative relationship between *N*_*e*_ and μ, although somehow expected theoretically (Sung et al.= 2012), is so far hypothetical.

Demographic fluctuations are another potential cause of discrepancy between genetic diversity-based and mutation load-based estimates of *N*_*e*_. Brandvain and Wright (2016) recalled that the mutation/selection/drift equilibrium is reached more quickly when selection is strong, with neutral mutations being the slowest to converge. This suggests that the mutation load might be less strongly influenced by ancient bottlenecks than the neutral genetic diversity, and therefore yield more reliable estimates of present-day drift power. In order to further investigate this hypothesis, we simulated coding sequence evolution in a population after a strong bottleneck. We found that the estimated 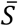 indeed equilibrates faster than π_S_ during the recovery phase (figure 4, generations 0-10,000). In these simulations, diversity-based estimates of *N*_*e*_ would be biased downwards during a substantial period of time after the bottleneck, whereas the mutation load would more quickly provide a reliable estimate. Ancient bottlenecks, therefore, might explain why genetic diversity-based and mutation load-based estimates of drift power sometimes disagree – *e.g.* see above our discussion of the *D. santomea* case. Can this effect account for the increased between species variance in drift power we report, compared to genetic diversity-based estimates? This would require additional assumptions, such as, *e.g.*, that large-*N*_*e*_ species tend to fluctuate more than small-*N*_*e*_ ones, or that the minimal value reached by *N*_*e*_ as populations fluctuate varies less among species than the maximal one. Such hypotheses have already been proposed (Romiguier et al. 2014) but so far lack any empirical support.

**Figure 4:**
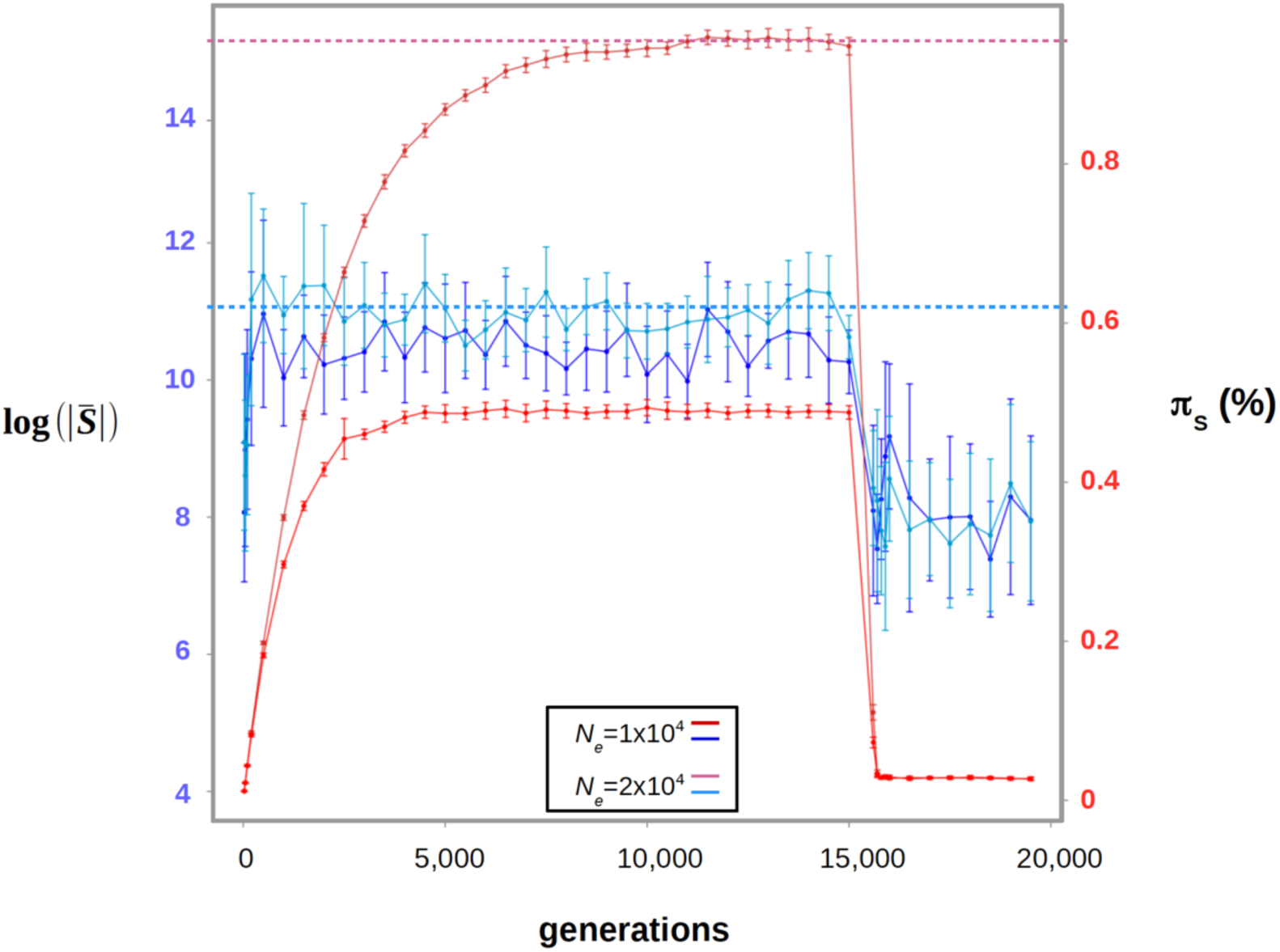
Simulation of the evolution of the estimated 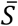 and π_S_ as *N*_*e*_ fluctuates. The simulation starts at time *t*=0 in a population devoid of polymorphism. The estimated 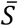 converges towards its equilibrium value faster than π_S_ during the recovery phase. Then a bottleneck is simulated at *t*=15,000 generations. The two statistics quickly reach their new equilibrium. Two population sizes were used: *N*_*e*_=10,000 and 20,000. Error bars represent the variance across 50 replicates. Simulated datasets in which the estimated shape parameter was smaller than 0.1 or greater than 0.6 were discarded – they yield extreme estimates of 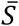. Horizontal dotted lines represent the equilibrium values of π_S_ and 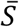 for *N*_*e*_=20,000.

A third and major factor potentially affecting the estimation of *N*_*e*_ is linked selection. The mutation load results from stochastic variations in allele frequency, which we have so far interpreted in terms of genetic drift. Linked selection – *i.e*., selective sweeps and background selection – is another source of stochasticity, which like drift is expected to result in a decreased genetic diversity and an increased mutation load (Kaiser and Charlesworth 2009, Barton 2010, Hartfield and Otto 2011). Corbett-Detig et al. (2015) demonstrated that the reduction in genetic diversity due to linked selection is stronger in large than in small population-sized species of plants and animals. Linked selection, therefore, tends to homogenize the genetic diversity among species. The impact of linked selection on the mutation load has been less thoroughly investigated, either theoretically or empirically. What we know is that recurrent selective sweeps result in patterns of neutral variation at linked loci that are best represented by a form of multiple-merger coalescent (Durrett and Schweinsberg 2005, Coop and Ralph 2012). Multiple-merger coalescents, on the other hand, are known to predict patterns of neutral and selected variation that depart the predictions of just drift (Eldon and Wakeley 2006, Der et al. 2012). So it might be that the respective effects of linked selection on the genetic diversity and the mutation load do not scale similarly with population size, perhaps explaining our results. This, again, is entirely hypothetical and would require to be confirmed via specific theoretical developments, which ideally should also account for the effect of background selection.

Finally, Castellano et al. (2019) recently introduced a new hypothesis. These authors reported a wider range of variation in 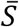 than in π among species of great apes, which they interpreted in terms of positive epistasis. Castellano et al. (2019) suggested that the average effect of a new deleterious mutation is negatively related to the existing load, because of the interaction between deleterious mutations. According to their interpretation, species genetic diversity would scale linearly with *N*_*e*_, whereas variations in 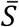 would reflect a positive correlation between *N*_*e*_ and 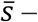 in contrast, we here assumed that 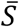 is constant across species. This interesting hypothesis offers yet another potential explanation to the discrepancy between diversity-based and mutation load-based estimators of drift power.

### 5. How variable among species is N_e_?

Lewontin’s paradox has been a recurrent cause of concern/excitement over the last decade (Leffler et al. 2012, Romiguier et al. 2014, Corbett-Detig et al. 2015, Coop 2016, Filatov 2019, Mackintosh et al. 2019). In animals, the within-species genetic diversity roughly spans two orders of magnitude, whereas population density and geographic range vary considerably more across species. Our analysis of the mutation load rather suggests that *N*_*e*_ varies by a factor of 10_3_, or maybe 10_4_, among species of animals. This is a step towards reconciling genetic with ecological estimates of population size – but how big is this step?

On one hand, one or two additional orders of magnitude can be seen as a moderate improvement, far from reconciling the effective and census population sizes of animal populations. Small insects or nematodes presumably outnumber large vertebrates by much more than a factor of 500 or 5000. On the other hand, Lewontin’s paradox may appear somewhat naive in suggesting that the genetic diversity should be proportional to the effective population size. Clearly, very large populations can just not follow the π=4*N*_*e*_μ prediction. This equation assumes mutation-drift equilibrium, which is only reached after a number of generations of the order of *N*_*e*_. As *N*_*e*_ increases, the assumption that the considered population has been devoid of bottlenecks and sweeps during the last ∼*N*_*e*_ generations becomes less and less plausible (Gillespie 2000). So maybe should we be satisfied, after all, by an estimated ratio of 10_3_ or 10_4_ of long-term *N*_*e*_ among species of animals? We suggest that Lewontin’s “paradox” in part reflects the varying definition/usage of the *N*_*e*_ parameter in the molecular evolutionary literature. Assessing the amount of stochasticity in allele frequency evolution, the prevalence of linked selection vs. drift, and their impact on genome evolution are key goals of current population genomics that perhaps do not need to be phrased in terms of a paradox, and probably cannot be reduced to just the issue of estimating one “*N*_*e*_” per species.

## Data accessibility

All the analyzed data sets are freely available from https://zenodo.org/record/3818299#.XramS-lS88o.

## Supplementary material

**Figure S1: Gamma model, separate analysis, primates vs. fruit flies, mean/shape correlation**

Blue: primates. Red: fruit flies. Both axes are in log scale.

**Figure S2: Gamma+lethal model, likelihood surface, primates + fruit flies.**

Legend: see figure 2.

**Figure S3: Relationship between π_n_/π_s_ and** 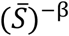**, primates + fruit flies.**

A linear relationship between the two plotted variables is theoretically expected at equilibrium with Gamma distributed effects (Welch et al. 2008).

**Figure S4: Gamma model, 47 species, shape parameter.**

Each dot is for a species. Y-axis: separate analysis: each species has its own shape parameter. X-axis: species from the same group share a common shape parameter.

**Figure S5: Gamma+lethal model, likelihood surface, 10 groups of animals**

Legend: see figure 2. X-axis: proportion of lethal non-synonymous mutations. Y-axis: Gamma shape parameter for non-lethal, non-synonymous mutations.

**Figure S6: Magnitude of variation in population scaled mean selection coefficient vs. neutral genetic diversity, Galtier 2016 data set.**

Each dot is for a species of animals. X-axis: relative synonymous heterozygosity. Y-axis: relative estimated 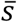. The highest two estimates of 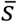 were obtained in nematode *Caenorhabditis brenneri* and mosquito *Culex pipiens.*

**Figure S7: Distribution of *r*_*i*_’s across species of primates (blue) and fruit flies (red)**

Model fitting was realized separately – no shared parameter among species. The Gamma+lethal model was applied to folded SFS, with parameter *p*_*lth*_ set to 0.6. Only one species of fruit flies has more than 9 categories of SNPs.

## Supporting information

responses to PCI comments

## Acknowledgements

We are grateful to Guillaume Achaz, Thomas Bataillon, Sylvain Glémin and Benoît Nabholz for insightful discussions. This work was supported by Agence Nationale de la Recherche grant no. ANR-15-CE12-0010. Version 3 of this preprint has been peer-reviewed and recommended by Peer Community In Evolutionary Biology (https://doi.org/10.24072/pci.evolbiol.100100).

