## Supplementary figures and images for "How much does *N*_*e*_ vary among species?"

### Supplemental Figure 1

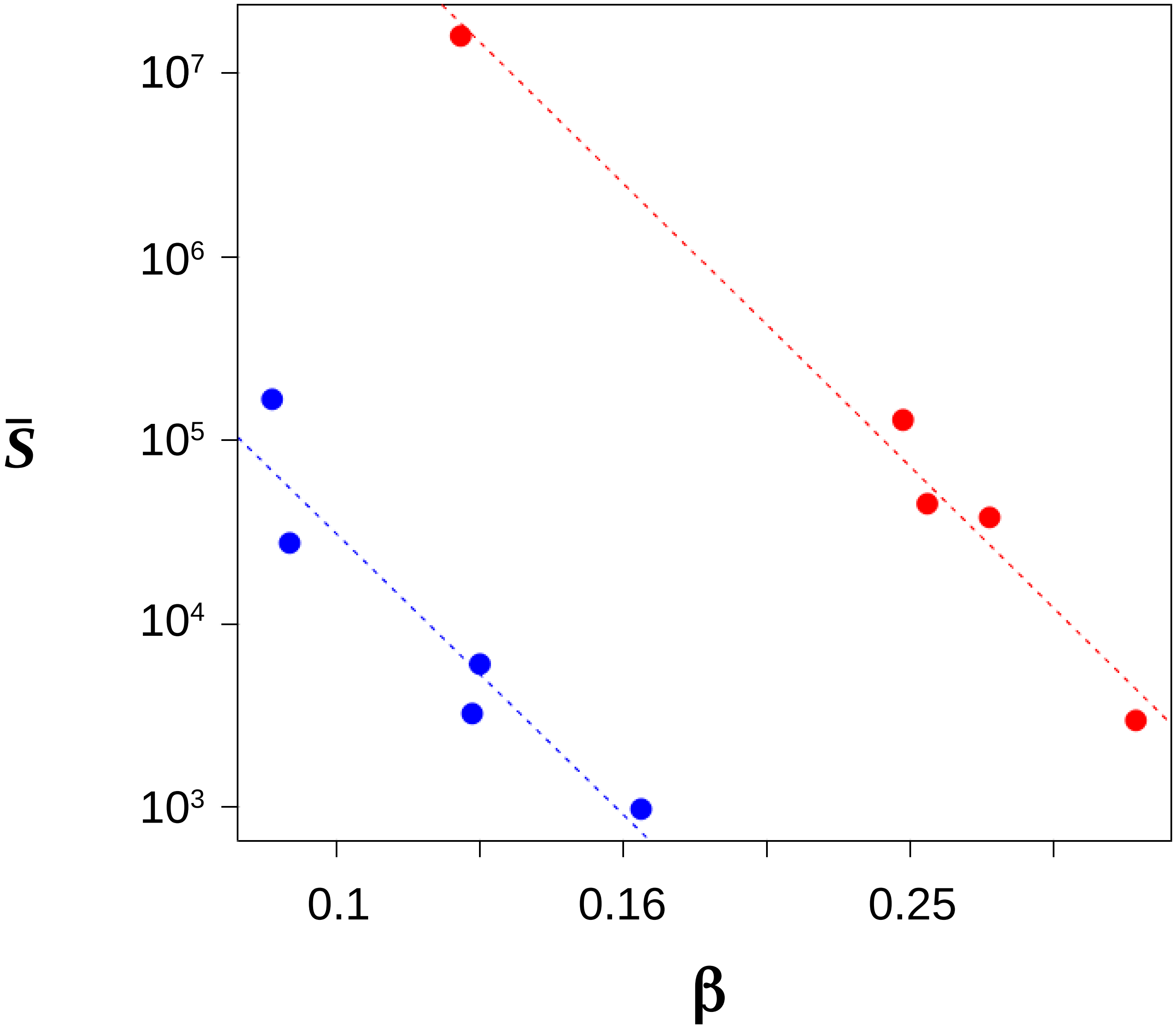

### Supplemental Figure 2

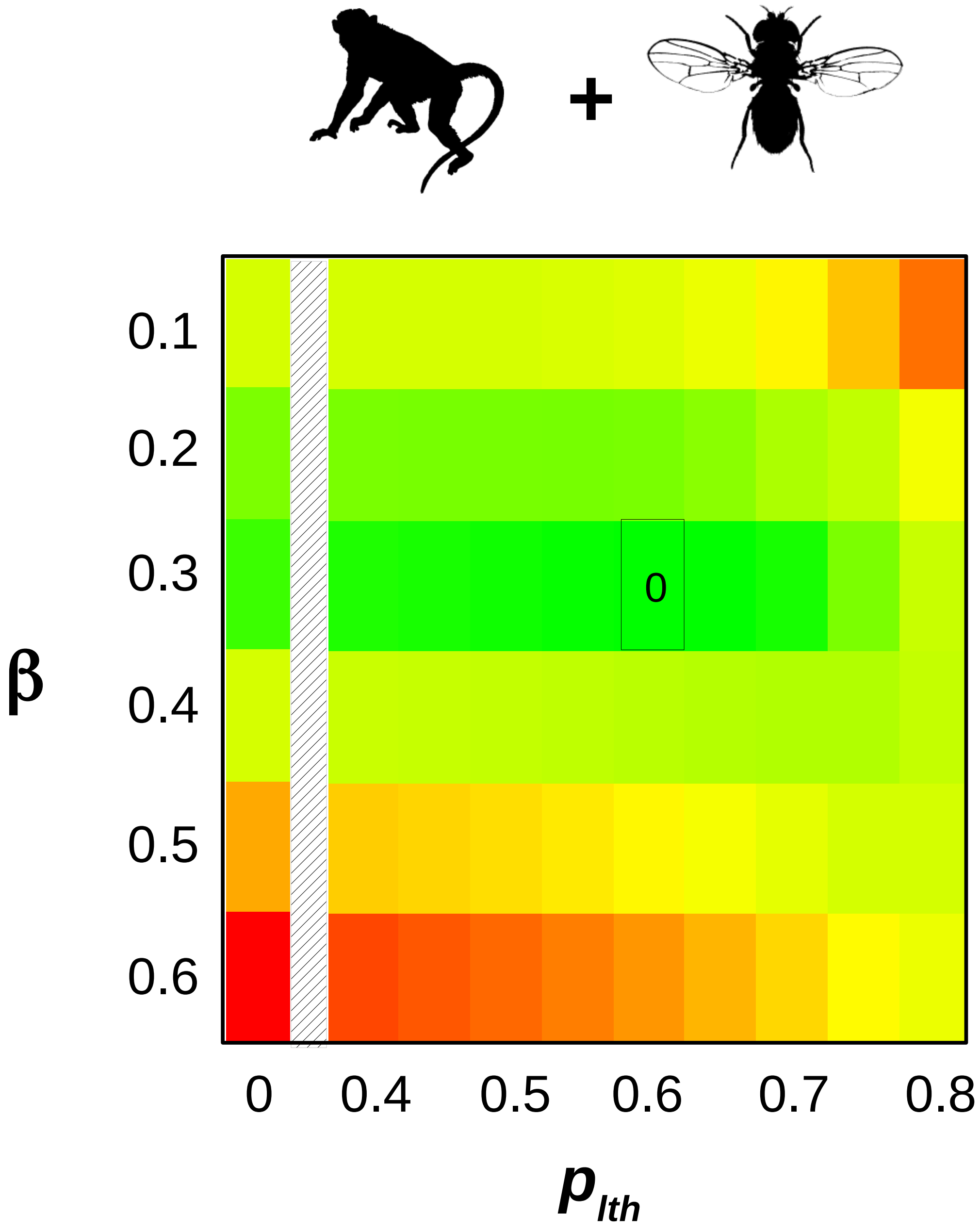

### Supplemental Figure 3

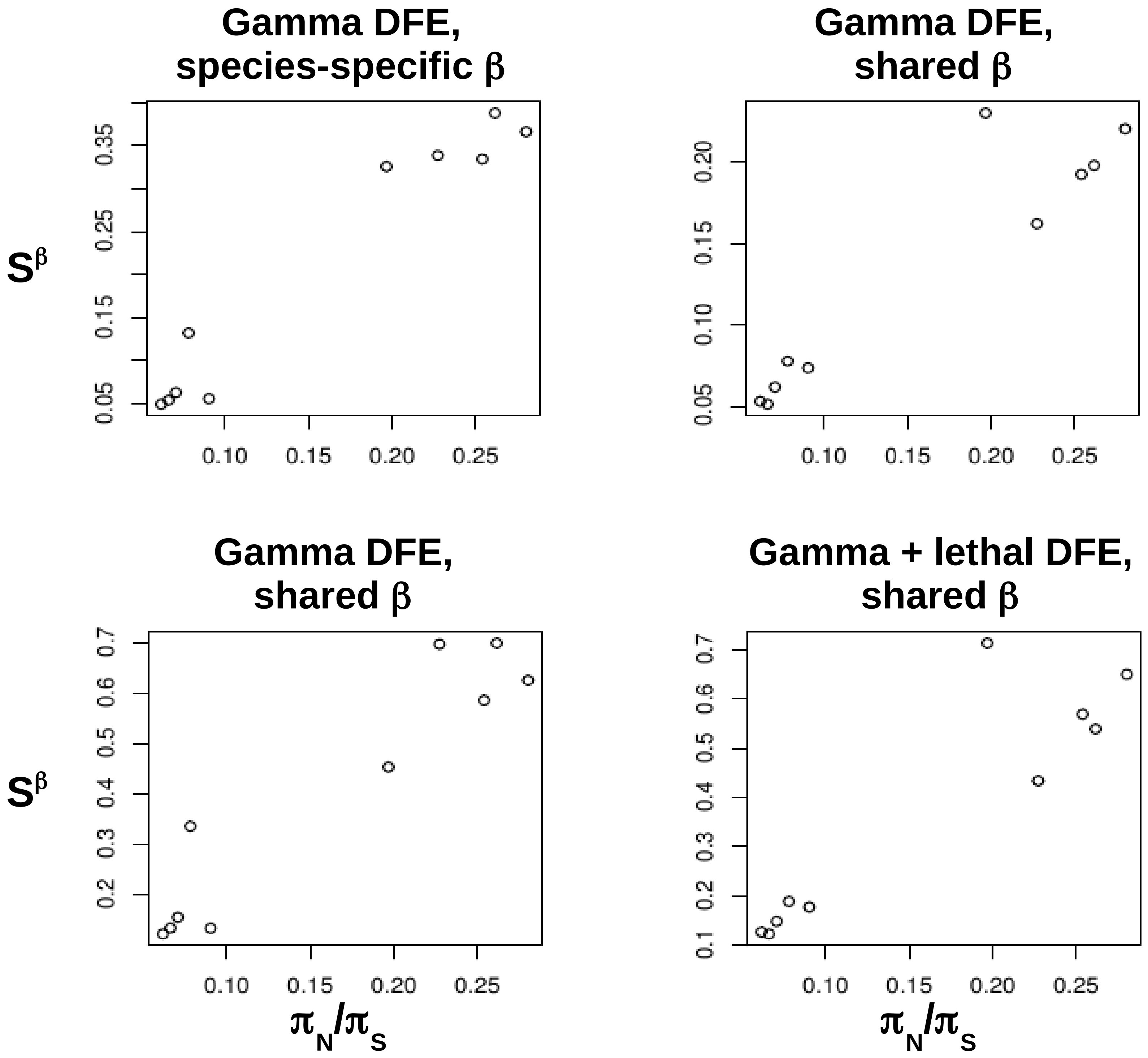

### Supplemental Figure 4

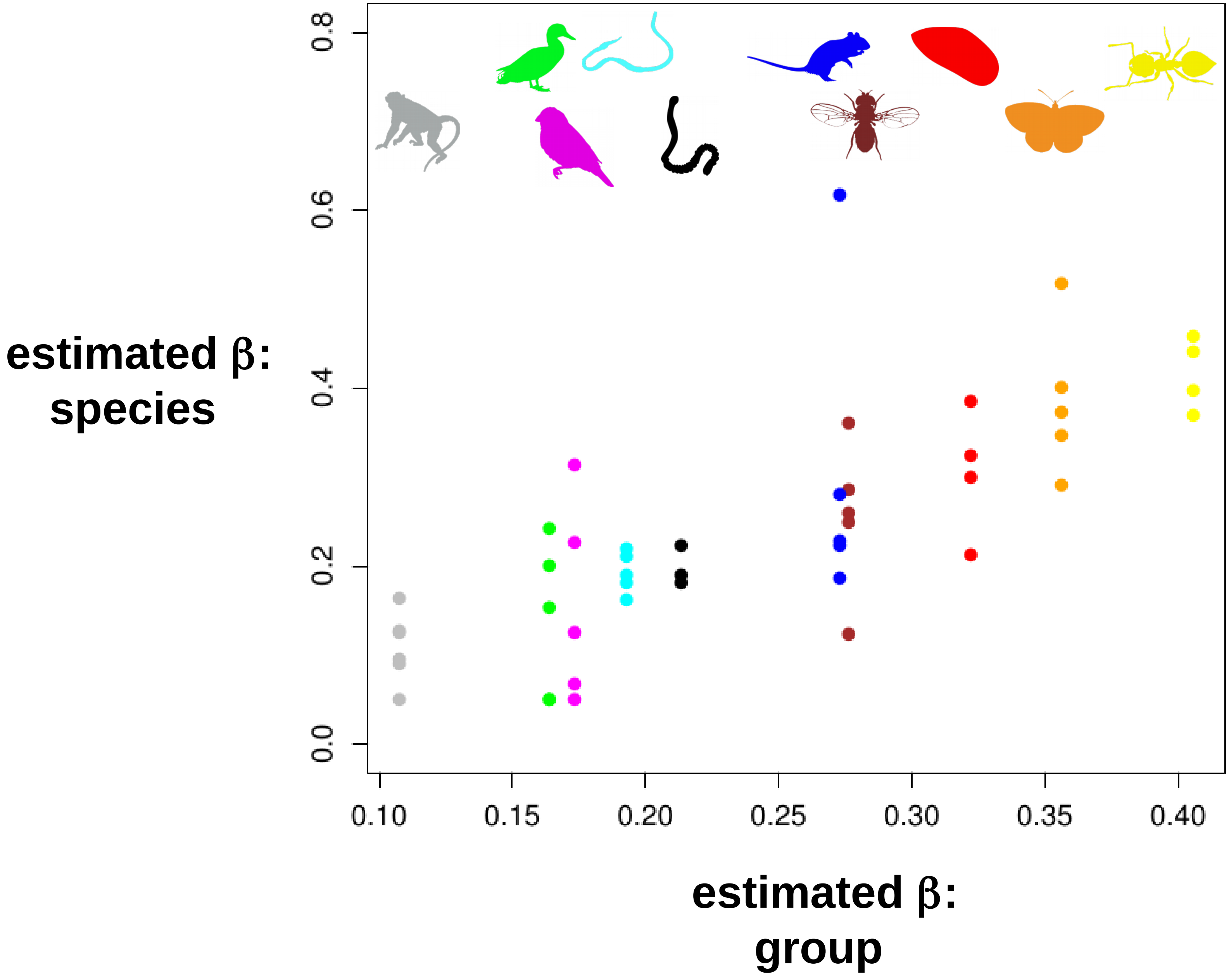

### Supplemental Figure 5

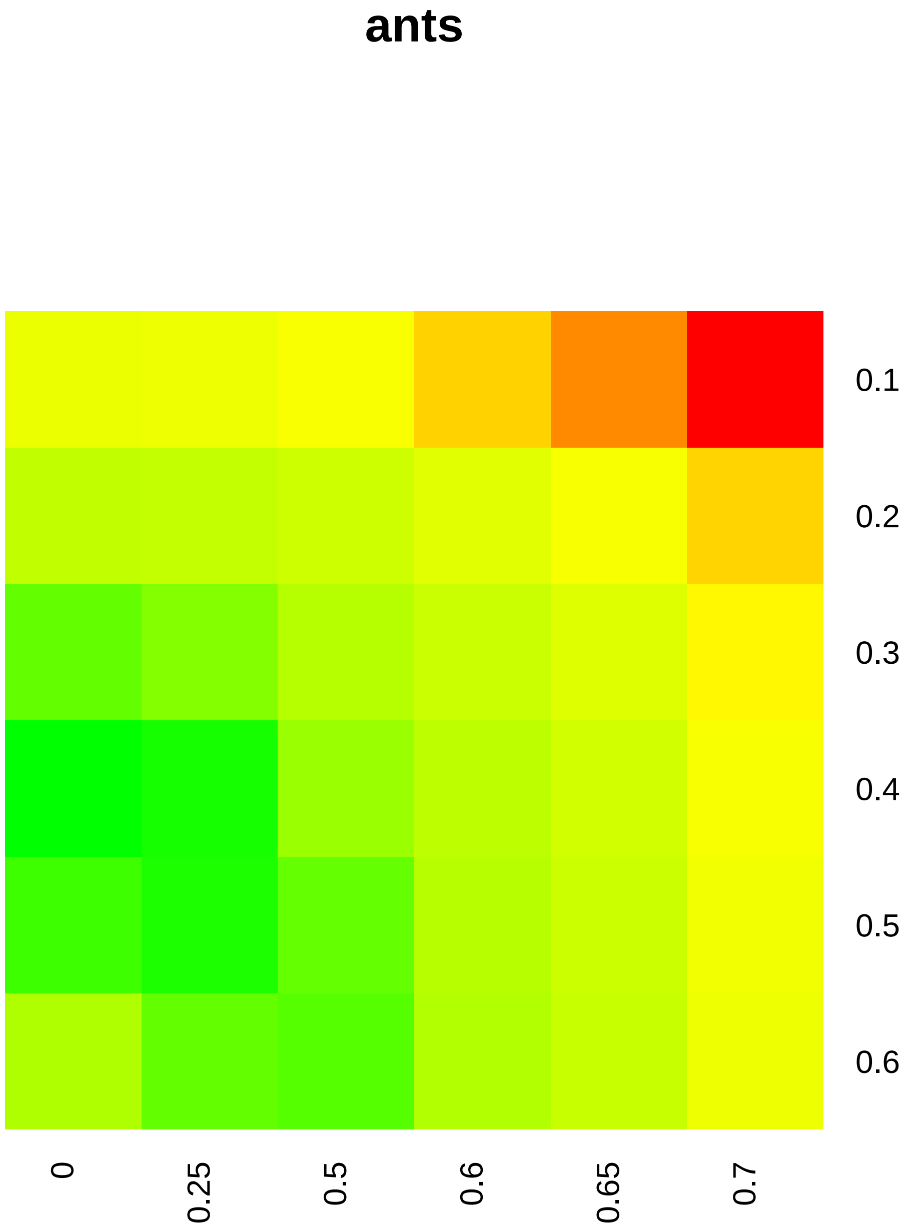

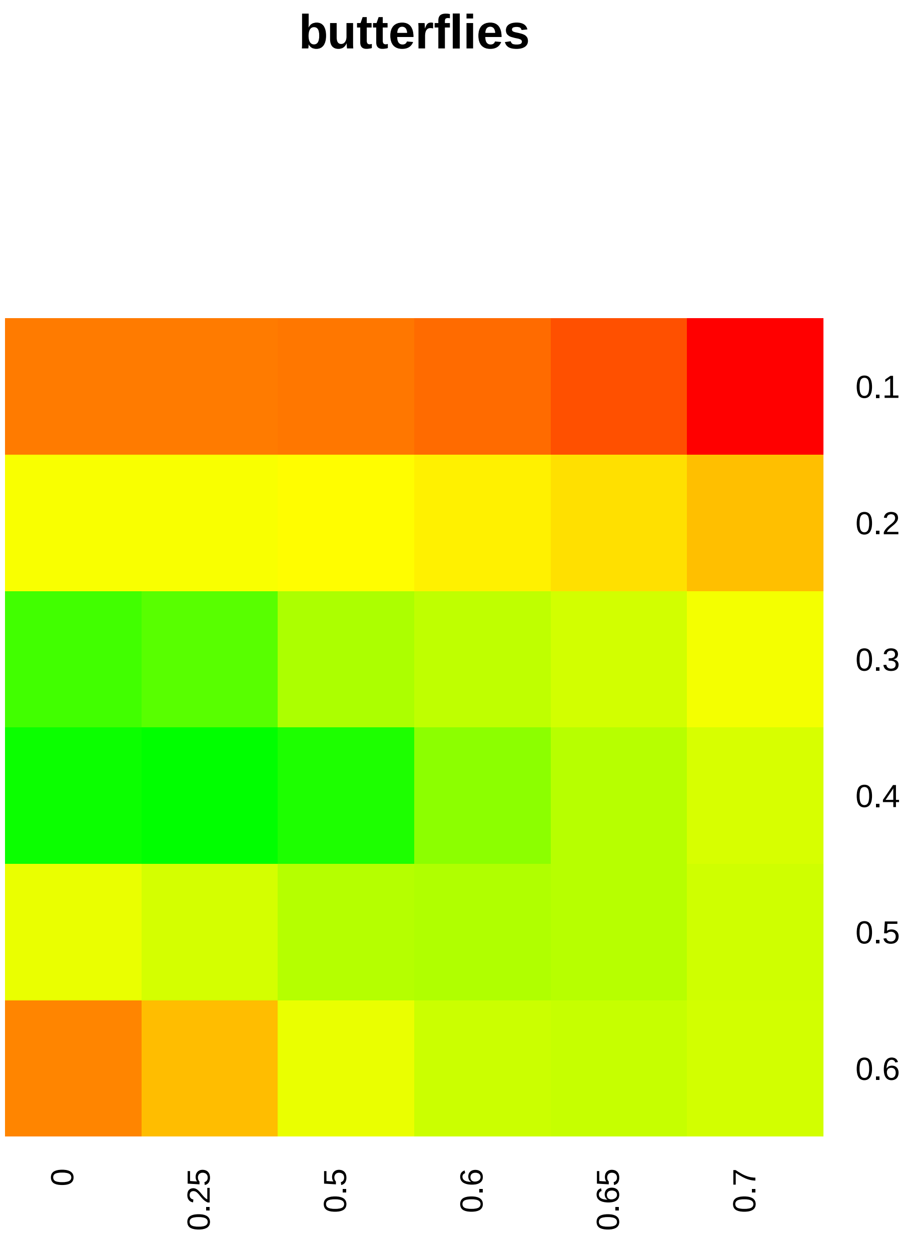

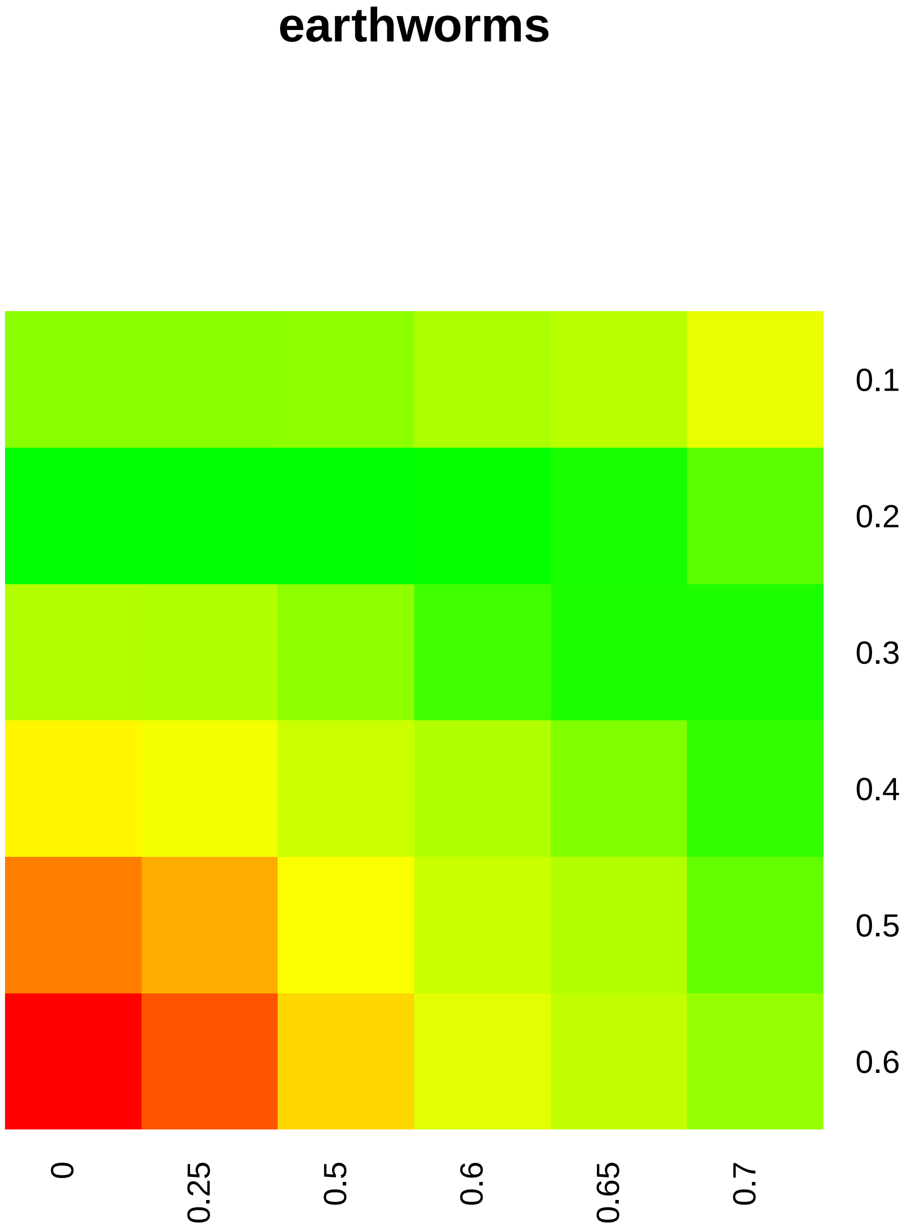

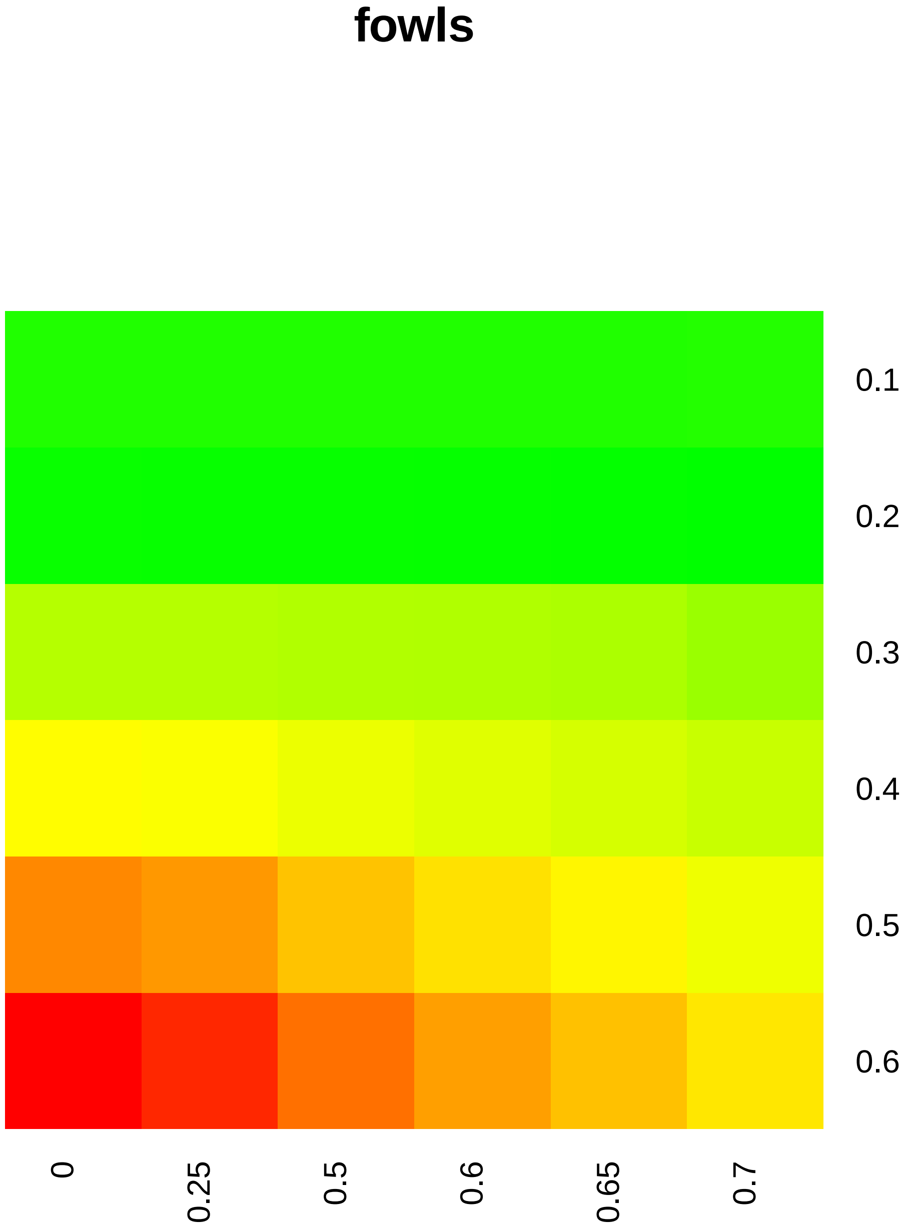

#### fruit flies

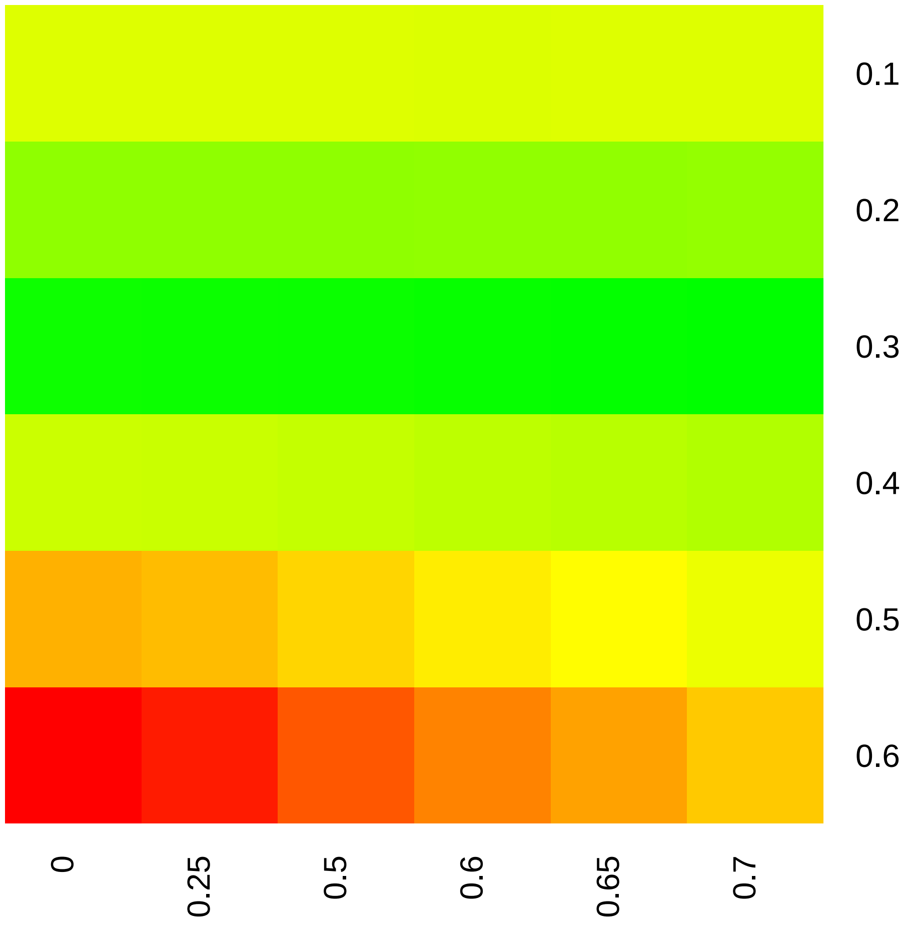

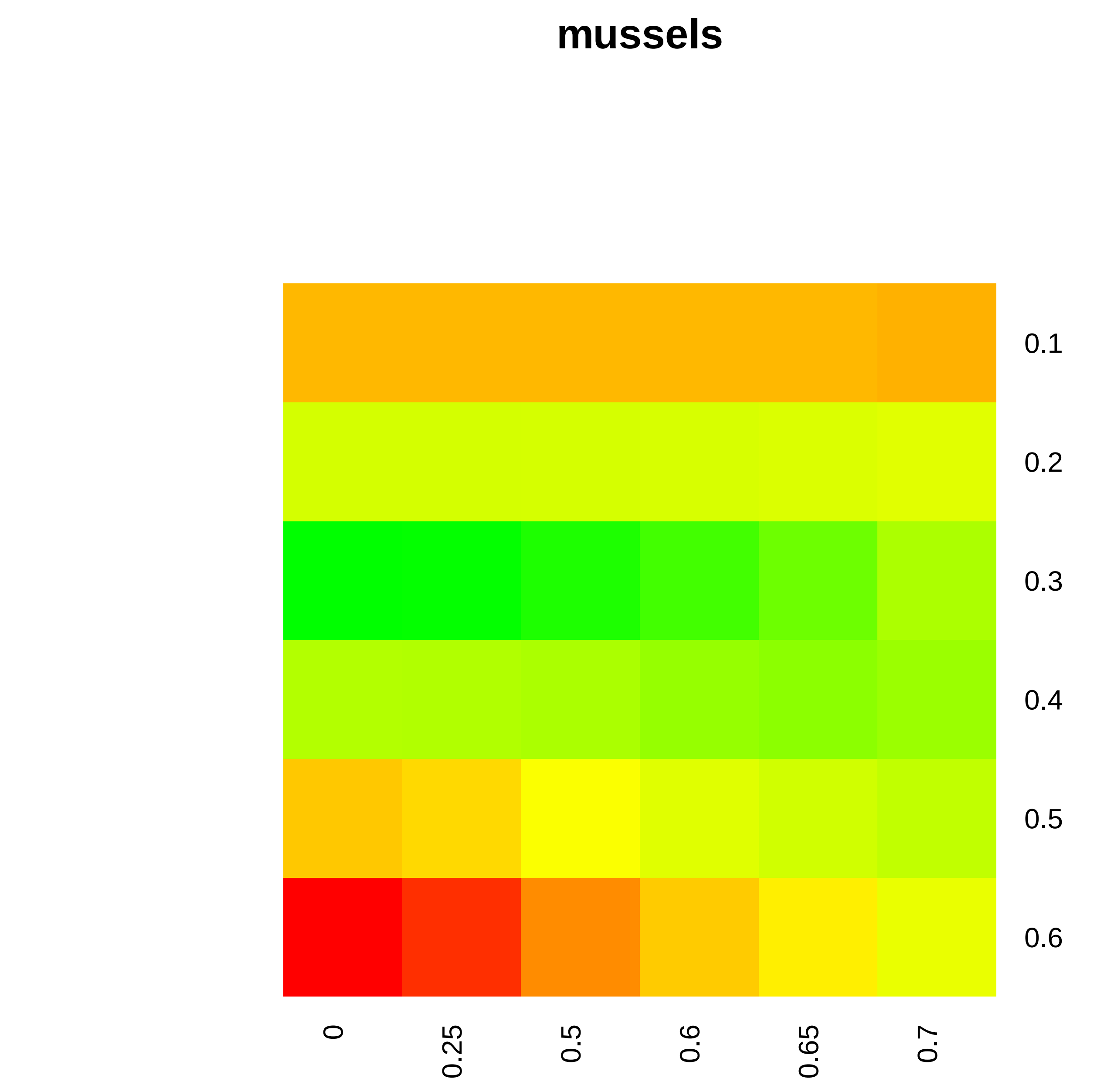

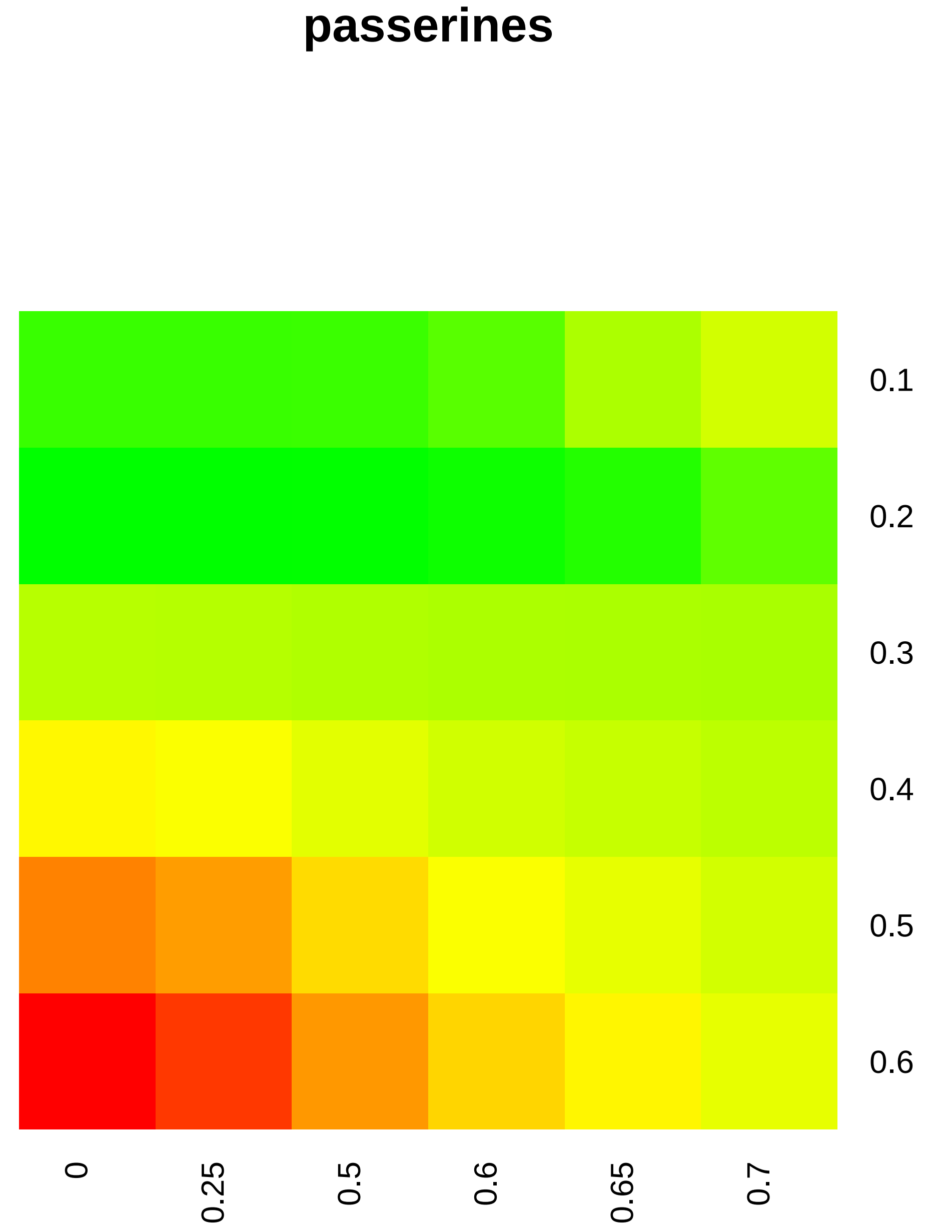

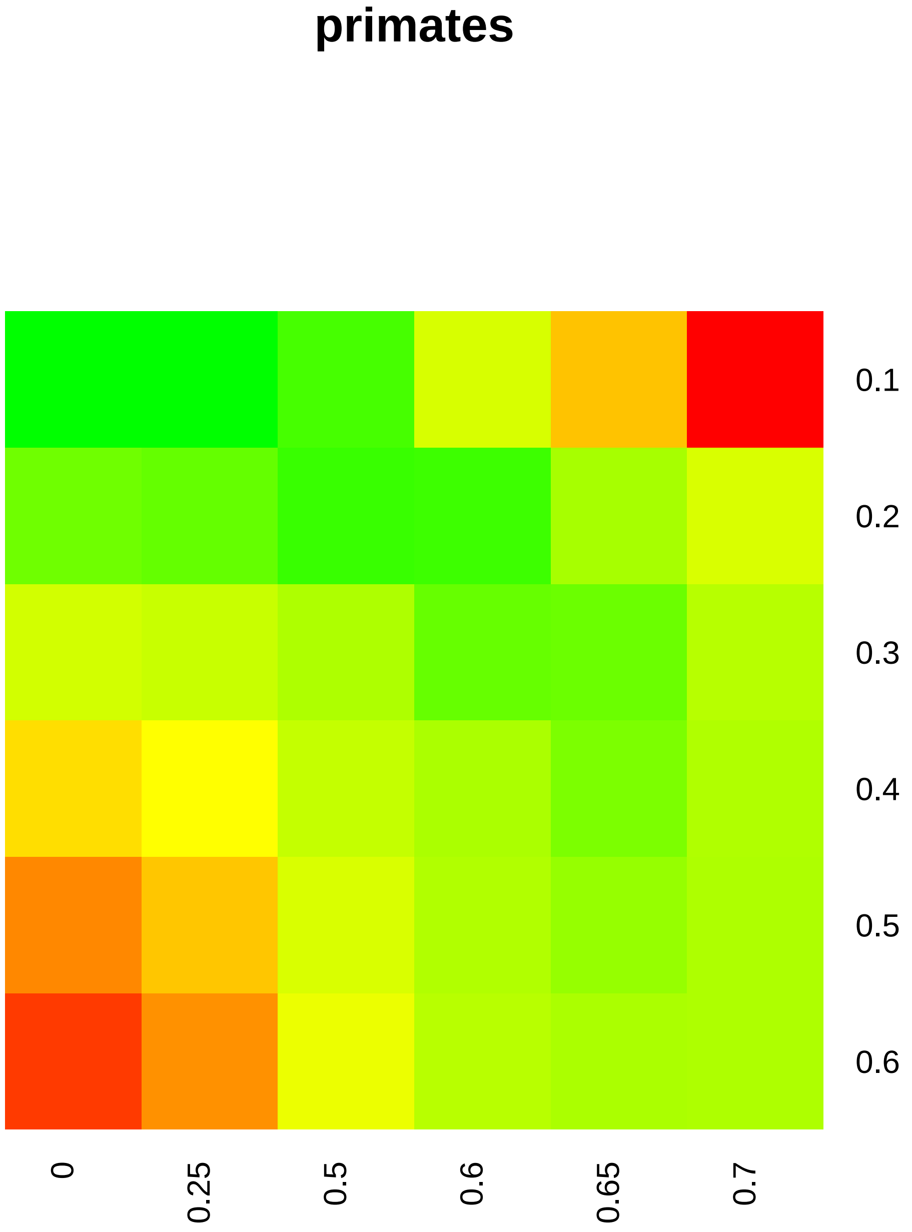

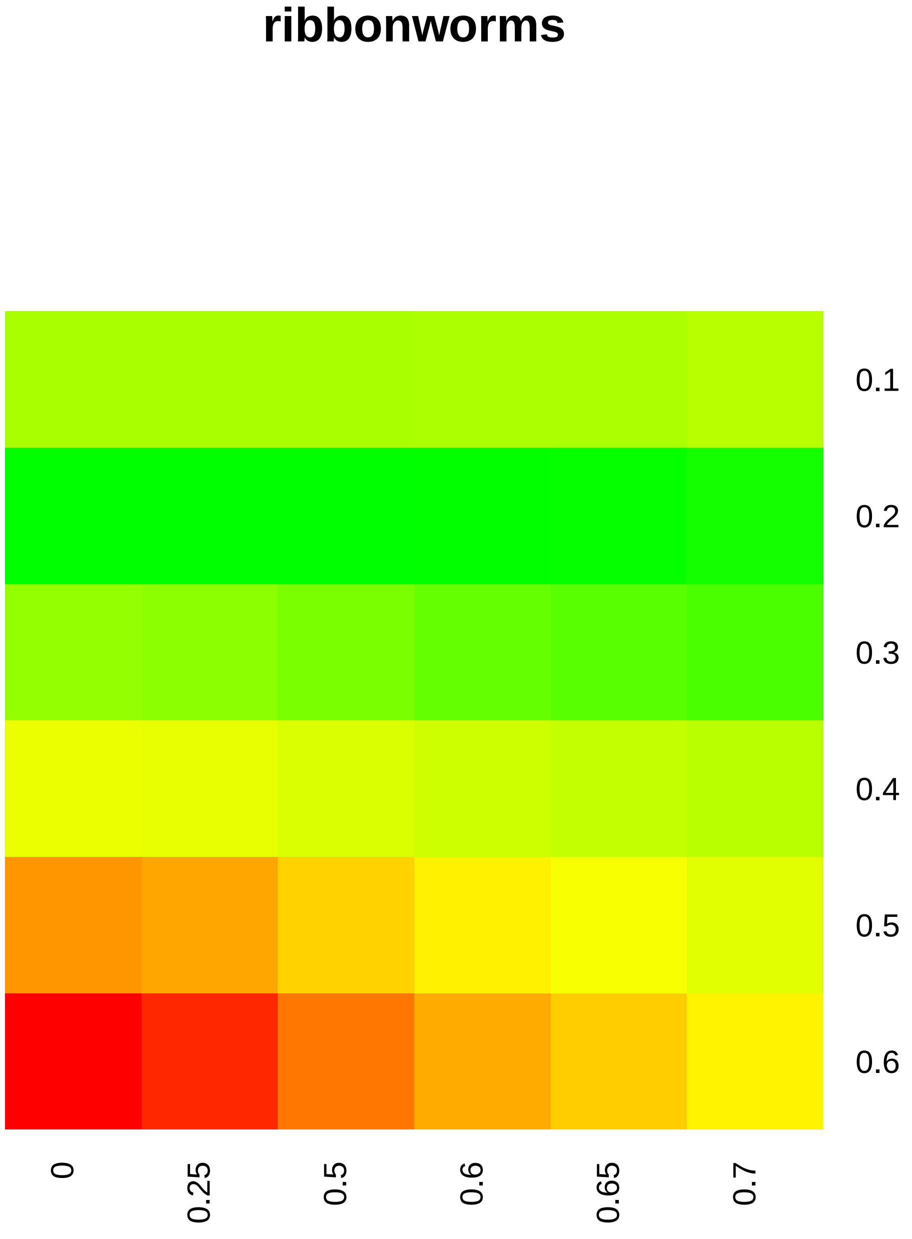

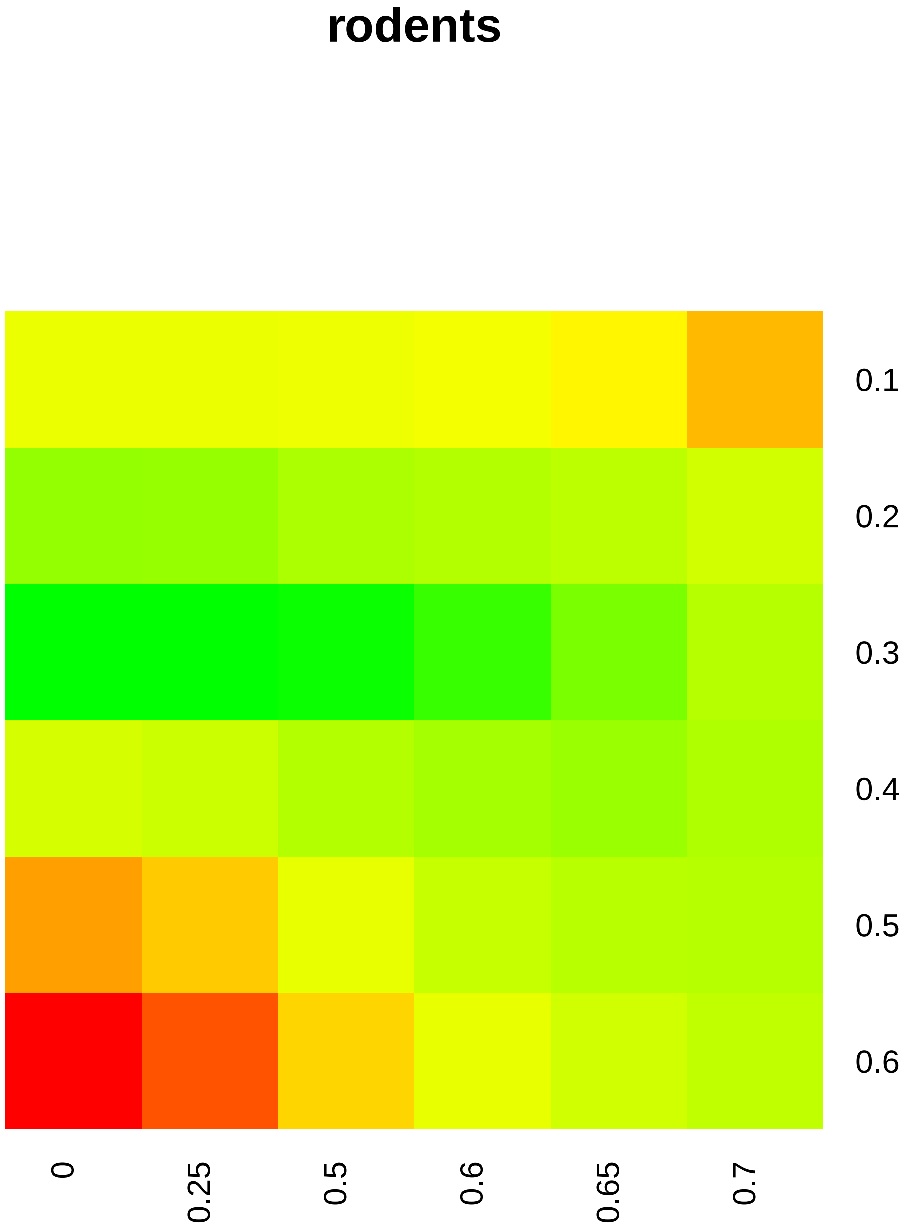

### Supplemental Figure 6

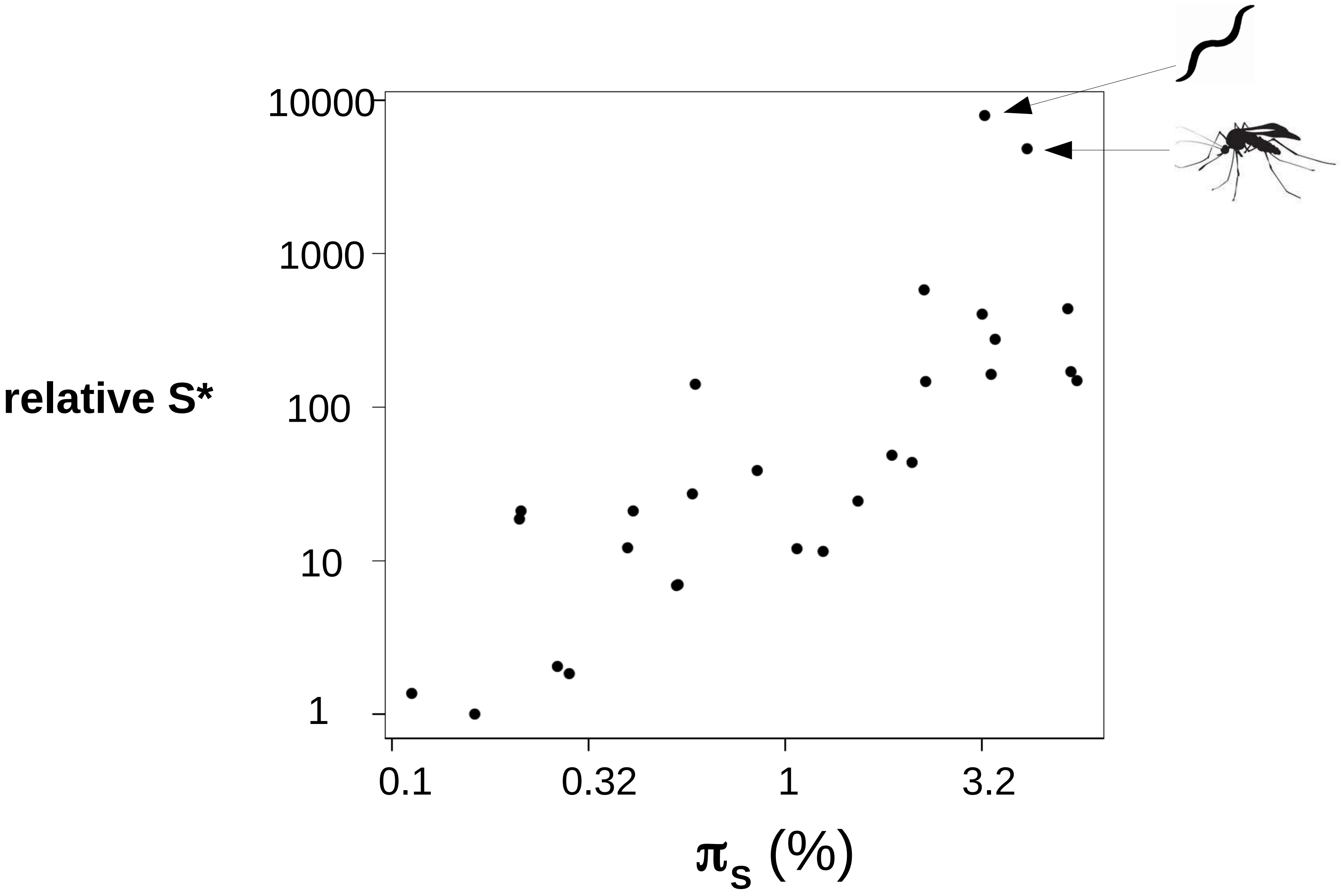

### Supplemental Figure 7

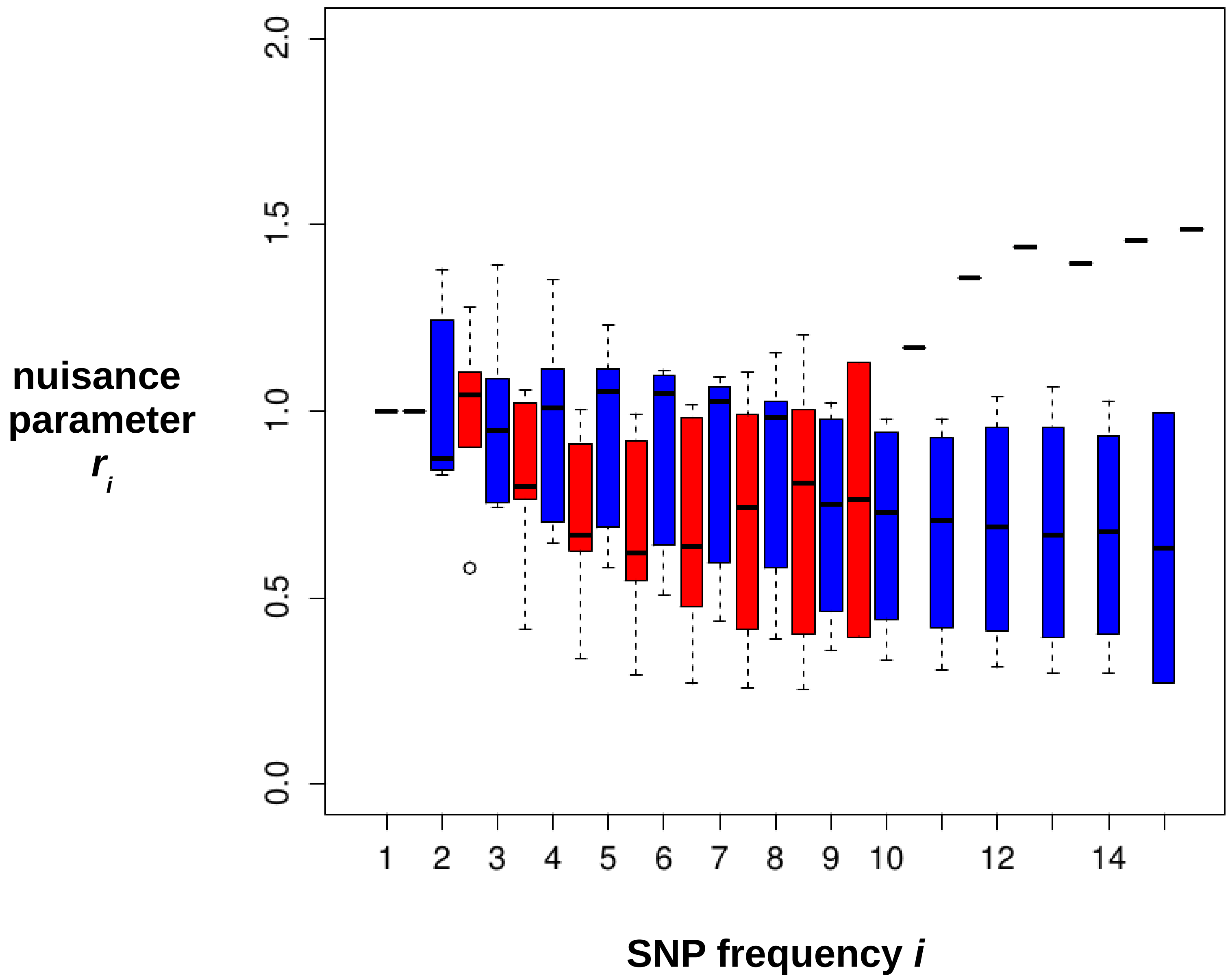
