## Supplementary material for "How much does *N*_*e*_ vary among species?": responses to PCI comments

**Dear PCI recommender and staff,**

**Thanks for this additional round of reviewing. We are sorry one of the reviewers got ill, and we wishes him/her a swift recovery. Please find below our response to the Reviewer 1's + Managing Board commants/recommendations of the. We hope  this version will be suitable for a recommendation by PCI Evol Biol.**

**Best regards**

**Marjolaine Rousselle, Nicolas Galtier**

Additional comments of the Managing board:

1) Mandatory modifications
As indicated in the 'How does it work?' section and in the code of conduct, please make sure that:

-Data are available to readers, either in the text or through an open data repository such as Zenodo (free), Dryad or some other institutional repository. Data must be reusable, thus metadata or accompanying text must carefully describe the data.

**The analyzed data (Site Frequency Spectrum, 101 species) are freely available from https://zenodo.org/record/3818299#.XramS-lS88o**
**as indicated in the preprint, l215-216. This archives includes documentation on the content of the data files and the command lines that were used to perform the analyses.**

-Details on quantitative analyses (e.g., data treatment and statistical scripts in R, bioinformatic pipeline scripts, etc.) and details concerning simulations (scripts, codes) are available to readers in the text, as appendices, or through an open data repository, such as Zenodo, Dryad or some other institutional repository. The scripts or codes must be carefully described so that they can be reused.

**The analyses have been performed using the multi_grapes program, which is freely distributed at https://github.com/BioPP/grapes**
**as indicated in the preprint, l232-233. The specific command lines that were used to perform the analyses are provided as part of the zenodo archive, see above**

-Details on experimental procedures are available to readers in the text or as appendices.

**All the analyses are described in the Material & Methods section and the zenodo archive.**

-Authors have no financial conflict of interest relating to the article. The article must contain a "Conflict of interest disclosure" paragraph before the reference section containing this sentence: "The authors of this preprint declare that they have no financial conflict of interest with the content of this article." If appropriate, this disclosure may be completed by a sentence indicating that some of the authors are PCI recommenders: “XXX is one of the PCI XXX recommenders.”

**Done**

2) when these changes are made, could you send us your MS in word or Latex format to. We'll try to format it according to PCI requirements.

**Done**

3) We will send it back to you for final verification and uploading to bioRxiv.

**Done**

Reviews
Reviewed by anonymous reviewer, 2020-04-09 12:29

I am grateful to the authors for taking my previous comments seriously. I think that the manuscript is much improved and remains interesting. Nevertheless, looking at the new Figure S7, I still have concerns about the legitimacy of estimating Ne from S_bar. If the authors agree, I think that the concerns should be emphasized more strongly. I also think that the authors could do more to show that the results could in principle be real. They need to show e.g. that a past bottleneck could lead to very large differences in the Ne values that apply to different statistics. Neither of these suggestions would require large changes.

Can Sbar be used to estimate Ne when the ri vary?

In the very welcome new section 460-474, the authors come close to stating that their key statistic, Sbar has no clear meaning.
They argue that, with highly variable ri, “what exactly \theta measures in this case is unclear”. I think \theta does have a clear meaning: the mutation rate multiplied by the length of the terminal branches in the genealogies. But the meaning of S_bar really is unclear, because it affects predictions for all frequency classes.

The authors then argue that the realized ~3-fold variation in the ri estimates (0.5-1.5; Figure S7) "does not suggest to us that the ri's pose a major problem of comparability in this analysis." This statement seemed too confident to me. Because the ri do not apply to different epochs, but to branches across the entirely genealogy, I think that the 3-fold differences imply quite major departures from a standard neutral genealogy, and it is not clear how Sbar will behave in those cases. For example, I think Figure S7 implies lots of high frequency polymorphisms in the flies, which is suggestive of balancing selection or structure.

Currently, it is suggested that the reader can skip sections 1-3 of the discussion, but, unless the authors think my comments above are mistaken, I think that sections 4-5 need to clearly acknowledge the possibility that S_bar does not provide a meaningful estimate of Ne.

**We generally agree with the author's comment. Whether the text is sufficiently cautious about this is, we suggest, a subjective matter. At any rate, the manuscript in its current form is more cautious that most, if not all, published McDonald-Kreitman-like analyses relying on $r_i$-based modeling. We would like to recall that our goal here is not to estimate S, but rather between-**

**species ratios of S. So if the potential problems raised by the reviewer apply more or less similarly to different species (which, we suggest, could be the case at least for the primates vs fruit flies comparison, see p 16), then we're essentially safe.**

**Following the reviewer's suggestion, we removed the sentence saying that the reader might skip the methodological discussion.**

Smaller comments/suggestions:

The simulations.

The simulations should prove to the reader that non-equilibrium demography could lead to very large differences in the range Ne values as estimated from piS and Sbar. Currently, this is not very explicit from Figure 4. Also, I could not find a description of the bottleneck depth at generation 15,000.

**We added in figure 4 horizontal lines representing the equilibrium vales of piS and S. Therefore the reader can appreciate that, during the first 1000 generations, a bottlenecked population would differ much from an equilibrium population in terms of piS, but not in terms of estimated S. We now clarify that the population size drops to Ne=500 at time 15,000 (legend to figure 4).**

The introduction

The introduction should probably acknowledge the classical results showing that populations can be characterized by different values of Ne when the Wright-Fisher assumptions are violated (e.g "inbreeding effective size" vs "variance effective size" etc.). These differences are not very surprising in themselves. What is surprising is the very large (~10-fold) differences estimated in this work.

**We agree but could not find a way to smoothly introduce this idea in the introduction.**

"To our knowledge, the Gamma + lethal model has been tried in two studies before this one." Didn't Nielsen and Yang 2003 MBE also use this model?

**Actually no, they did not. Nielsen & Yang used a wide range of model, including the Reflected Gamma and Reflected Gamma + lethal, but not the Gamma + lethal. This analysis relied on divergence data, not polymorphism data, a limited number of genes, and does not seem directly comparable to ours. For instance, the estimated shape parameter of the reflected Gamma was >3 in Nielsen & Yang 2003, whereas polymorphism-based estimates are almost always well below 1.**

In my previous comment 2.3, I asked whether the authors might attempt to estimate s_bar (i.e the mean selection coefficient unscaled by Ne). I think that previous authors have done this, and it relates to the interesting discussion in lines 631-635.

**To estimate s_bar = S_bar/(4$N_e$), one needs an estimate of 4$N_e$. Chen et al (2017) used $\pi_4/\mu$ as an estimate of $N_e$, where $\pi_4$ is the heterozygosity at four-fold positions (very similar to our $\pi_S$) and $\mu$ is an estimate of the per generation mutation rate. They found that the estimated s_bar varied extensively between species, and doubted its reliability. By estimating $N_e$ from the heterozygosity at neutral sites, the Chen et al (2017) approach implicitly assumes that $\theta$ and S_bar are proportional, whereas our study rather focuses on the discrepancy between these two statistics**

**(e.g. see section 4 of the discussion). For this reason we refrained to do this calculation in the current study, since it relies on an assumption that we question in the first place.**

What are the CIs in Figure S7?

**Each vertical box in figure S7 represents the distribution of one estimated $r_i$ among the five species of primates (blue) or fruit flies (red) we analyze.**

Is Kaiser and Charlesworth 2009 TIG relevant to the discussion of linked selection and mutation load? (lines 621-625).

**Very relevant, thanks for mentioning this, now cited.**

l445: “can be interpreted as the relative effective mutation rate”. Could r_i be more clearly described as the relative length of the portions of the genealogies that lead to i tips?

**We are not sure this description is entirely correct, or one needs to add "divided by the expectation assuming demographic equilibrium". The expected number of synonymous SNPs at frequency $i$ is equal to $r_i/i$, not just $r_i$ (with $1/i$ being the Wright-Fisher expectation), and similarly for the non-synonymous SFS (see equations 6 in 7 in Galtier 2016).**

There are two different legends for Figure 4 (pp. 35 and 39).

**Thank you for spotting this, now corrected.**

Grammar etc.
l38: "which implies to also infer" should be "which implies also inferring".
l51: "distribution of fitness effect[s]".
l72: "can as well be" should be "can also be".
l101: "by jointly analysis" should be "by jointly analyzing".
l242 and Figure 4. Does 2.10^4 mean 20,000? If so, notation could be clearer.
l243: 2.2 10^{-7} (missing multiplication sign).
l269: "we rather fitted" should be "we instead fitted"
l303 and l310: "probability to be observed" should be "probability of being observed"
l331: "Setting plth [set]".
l338: "colons" should be "columns".
l440: “parameters of nuisance” should be “nuisance parameters”.
l464: “in [the] absence of”.
l536: “implies to accommodate” should be “implies accommodating”.
l556: “did not change much the picture” should be “did not change the picture much”.

**All of these corrected, thanks.**
